# Neoplastic immune mimicry potentiates breast tumor progression

**DOI:** 10.1101/2025.01.17.633673

**Authors:** Eric B. Berens, Sokchea Khou, Elaine Huang, Amber Hoffman, Briana Johnson, Nell Kirchberger, Shamilene Sivagnanam, Nicholas L. Calistri, Daniel Derrick, Tiera A. Liby, Ian C. McLean, Aryn A. Alanizi, Furkan Ozmen, Tugba Y. Ozmen, Gordon B. Mills, E. Shelley Hwang, Pepper J. Schedin, Hugo Gonzalez, Zena Werb, Laura M. Heiser, Lisa M. Coussens

## Abstract

Dedifferentiation programs are commonly enacted during breast cancer progression to enhance tumor cell fitness. Increased cellular plasticity within the neoplastic compartment of tumors correlates with disease aggressiveness, often culminating in greater resistance to cytotoxic therapies or augmented metastatic potential. Here we report that subpopulations of dedifferentiated neoplastic breast epithelial cells express canonical leukocyte cell surface receptor proteins and have thus named this cellular program “immune mimicry.” We document neoplastic cells engaging in immune mimicry within public human breast tumor single-cell RNA-seq datasets, histopathological breast tumor specimens, breast cancer cell lines, as well as in murine transgenic and cell line-derived mammary cancer models. Immune-mimicked neoplastic cells harbor hallmarks of dedifferentiation and are enriched in treatment-resistant and high-grade breast tumors. We corroborated these observations in aggressive breast cancer cell lines where anti-proliferative cytotoxic chemotherapies drove epithelial cells toward immune mimicry. Moreover, in subsequent proof-of-concept studies, we demonstrate that expression of the CD69 leukocyte activation protein by neoplastic cells confers a proliferative advantage that facilitates early tumor growth and therefore conclude that neoplastic breast epithelial cells upregulating leukocyte surface receptors potentiate malignancy. Moving forward, neoplastic immune mimicry should be evaluated for prognostic utility in additional breast cancer cohorts to determine its potential for patient stratification. Future research should evaluate correlates with distal metastases, progression-free survival, overall survival, and therapeutic response/resistance.

**Statement of Significance:** Neoplastic breast epithelial cells express surface receptors canonically attributed to leukocytes and are associated with therapy resistance and aggressive tumor behavior.

## Introduction

Tumors are complex ecosystems containing a variety of interacting cell types and cell states (1). Research has sought to identify the cells driving tumor progression and this effort has led to both epithelial and stromal candidates (2,3). The search for potentiators of malignancy has produced seminal discoveries including dedifferentiated tumor-initiating cells that fuel tumors despite existing in low abundance (4). Aggressive tumors driven by dedifferentiated subpopulations also exhibit therapeutic resistance programs along with increased tumor recurrence and metastatic potential (5,6). The plasticity of dedifferentiated neoplastic cells renders them difficult to clinically locate and eradicate, potentially contributing to cancer patient mortality (7,8). Unveiling the biomarkers and functions inherent in aggressive neoplastic cells could underlie more effective therapies and thereby improve cancer patient outcomes. These are pressing tasks as the global burden of cancer continues to rise (9).

Neoplastic cells that acquire stem-like characteristics often gain the ability to undergo metaplasia or mimic various stromal cell types (10,11). Studies have demonstrated that primary tumor growth and metastasis are fostered by neoplastic cells harboring transcriptional programs reminiscent of fibroblasts (12), endothelia (13), or even neurons (14). The notion that neoplastic cells might also mimic leukocytes is suggested by recent research documenting unexpected expression of surface markers canonically attributed to leukocytes (15–22). Highlights of this involve aggressive breast carcinoma cells expressing CD14, a prominent myeloid protein also reported on breast epithelia (15), stem-like cutaneous squamous carcinoma cells expressing CD80 that typifies B cells (16), and pancreatic adenocarcinoma cells expressing the myelomonocytic indicator CD68 (17). There is additional evidence that neoplastic epithelial cells upregulate the CD74 invariant chain of the MHC II molecule (18) or lymphocyte chemokine receptors, including CXCR4 when disseminating (19). Neoplastic cells can also acquire expression of leukocyte CD45 via fusion with myeloid cells to generate circulating hybrids that promote metastasis (20). In each of these instances, neoplastic cells appear to utilize canonical leukocyte surface receptors to enhance their intrinsic malignancy or manipulate their stromal microenvironments.

In this study, we identified subpopulations of dedifferentiated neoplastic epithelial cells expressing biological programs ascribed to endothelia, mesenchyme, neurons, or leukocytes in a single-cell RNA-sequencing (scRNA-seq) analysis of 70 human public breast tumors (23,24). We focused further on “immune mimicry” as an emergent phenotype accompanying aggressive treatment-resistant disease. Immune-mimicked neoplastic cells upregulate variable combinations of ∼20 surface marker genes canonically expressed by leukocytes including CD3, CD14, CD18, CD45, CD68, CD69, CD74, and CD83, which were found in nearly every breast tumor specimen. We corroborated this cell state in 12 human breast and murine mammary cancer cell lines, where cytotoxic therapy further induced immune mimicry in aggressive/metastatic lines. Neoplastic cells expressing the CD69 leukocyte activation marker (25,26) were additionally validated in a cohort of 58 histopathological breast tumor specimens via multiplexed immunohistochemistry. We subsequently linked neoplastic CD69 to enhanced tumor growth in experimental mammary cancer models. Together, our observations indicate that subsets of aggressive neoplastic breast epithelial cells express canonical leukocyte proteins, a molecular program that appears widespread in breast cancer.

## Materials and Methods

### Study Design

This study sought to provide evidence that small subpopulations of epithelial breast cancer cells express canonical leukocyte surface receptors and that this “immune-mimicked” cell state is associated with malignancy. In endeavoring to achieve this objective, we conducted scRNA-seq and histopathological analyses on primary human breast tumors, further supporting our findings with experimental studies using related cell lines and mice. Of note, sample sizes were not determined in advance, instead, for our scRNA-seq and histopathological analyses, they were foremost dictated by the availability of datasets and resources. For cell line experiments, three to five biological replicates were generally chosen to adequately portray variability inherent within the reported phenotypes. Mouse experiments were also guided by resource availability for which a variety of distinct models were employed to build confidence in the results. All mouse experiments were approved by our Institutional Animal Care and Use Committee. Reagent tables referenced in the Materials and Methods section include Resource Research Identifiers (RRIDs).

### scRNA-seq analysis

Count matrices for human breast tumor scRNA-seq datasets (23,24) were downloaded from the Gene Expression Omnibus (GSE161529, GSE176078) and supplemented with additional tumors from the PANNTHR trial at the Oregon Health & Science University (Table S1). Count matrices for reduction mammoplasties (27) were also downloaded from the Gene Expression Omnibus (GSE235326). All scRNA-seq analyses were conducted in R using Seurat-V5 (28). Tumor samples containing all cells were first processed separately and subjected to filtering for nfeatures (lower limit >200; upper limit tailored per sample) and mitochondrial reads (<10% for Pal *et al.* and PANNTHR (NCT04481113) datasets, < 20% for Wu *et al.* dataset). DoubletFinder (29) was then applied to each sample to identify and remove cellular multiplets, assuming a ∼5% multiplet rate per ∼10,000 cells. Singlet cells were next collected for downstream analysis after which epithelia were extracted using a gating threshold > 1 for established mammary cytokeratin biomarkers KRT14, KRT18, or KRT19. Epithelial cells were extracted from the Kumar *et al.* reduction mammoplasty dataset (27) in the manner described above. InferCNV (RRID:SCR_021140) was subsequently performed on individual tumor samples to identify neoplastic epithelium by comparison to an epithelial reference. This reference was built by combining 100 epithelial cells downsampled from the 124 reduction mammoplasty samples in which we ascertained 8 reference groups. We considered tumor epithelial cells to be neoplastic if chromosomal CNVs were inferred versus these 8 reference groups.

After identifying neoplastic epithelia in each tumor sample, or mammary epithelia in the reduction mammoplasties, we next merged scRNA-seq samples within datasets. Our decision to merge (i.e. not integrate) samples without regressing cell cycle status was predicated on a desire to maintain interpatient heterogeneity. Diverse scRNA-seq clusters were identified using Shannon Weaver Diversity Index scoring via the vegan R package (RRID:SCR_011950) and clusters were considered diverse if this score was above 1.5. Genes significantly upregulated in diverse clusters were subjected to FGSEA (RRID:SCR_020938) to identify significantly enriched Gene Ontology Biological Processes (GO-BPs) obtained via MSigDB (30). Neoplastic (or epithelial) cells were defined as immune-mimicked if they resided within a diverse scRNA-seq cluster with enrichment of immune-related GO-BPs. Immune-mimicked clusters were further given myeloid-like or lymphoid-like designations based on enrichment of myeloid- or lymphoid-related GO-BPs. Surface receptors attributed to myeloid-like and lymphoid-like phenotypes involved filtering differentially enriched genes with a list of human cluster of differentiation (CD) surface markers from UniProt (release: 2024_06). CD receptors were only considered if they showed a fold-change increase of >1.5 and p <0.001 in immune-mimicked scRNA-seq clusters. When comparing inferred CNVs in CD45-pos versus CD45-neg neoplastic cells, we extracted CNV calls from inferCNV HMM predictions, where states :: 2 were classified as deletions and ≥ 4 as amplifications. We then compared the genes and genomic regions (coordinates) comprising specific inferred CNVs, later aggregating the number of CNVs per chromosome to create subpopulation profiles that could be correlated across patients. These analyses only considered patients with at least 10 CD45-pos neoplastic cells for statistical reasons.

Many scRNA-seq plots presented in the manuscript were generated by quantifying either the percent of neoplastic/epithelial cells engaging in immune mimicry or the percent of cells expressing a given receptor. These positivity comparisons were done to facilitate drawing parallels between scRNA-seq and flow cytometric analyses. Although sample merging was done at the dataset level during scRNA-seq analysis, all samples were combined when making plots with extracted expression matrices, linked metadata, and clinical attributes. Signatures used for comparing immune-mimicked and non-mimicked cells were also obtained from MSigDB (30), the cell cycle analysis followed the Seurat tool’s vignette, and the correlation plot was generated with R’s corrplot package (RRID:SCR_023081). Last, the human PBMC dataset used for scRNA-seq analysis was accessed via the portal accompanying its publication (31).

### Multiplexed immunohistochemistry

Primary breast tumor tissue was acquired from two sources for our histopathological evaluation of immune mimicry, notably epithelial CD69. These sources encompassed a breast tumor tissue microarray created from 26 patient specimens retrospectively banked at the Oregon Health & Science University’s Knight Cancer Institute BioLibrary, which we supplemented with 32 other breast tumor tissues from a previous cohort (32). FFPE tissue sections of ∼5 μm thickness from this pooled cohort of 58 primary breast tumors were then subjected to multiplexed immunohistochemistry (mIHC) as previously described (33,34). The antibodies we used are found in Table S2 and, notably, the CD69 antibody was specifically chosen due to its widespread use in The Human Protein Atlas: (https://www.proteinatlas.org/) (35) . To detect surface receptor protein expression via mIHC analysis, we first created a nuclei mask from the H3NUCA and then expanded the mask based on PanCK staining using a propagation algorithm in CellProfiler (36) to obtain whole epithelial cell segmentation. Each cell’s segmentation was expanded if it had PanCK stain while neighboring cells maintained their original nuclei segmentation. The mean intensity of each marker was measured per cell and characterized as positive or negative contingent on visually validated thresholding in the Image Cytometry version of FCS Express. See protocol for additional details: dx.doi.org/10.17504/protocols.io.n92ldmmznl5b/v2. Epithelial CD69 was distinguished in the with the following cellular gates: PanCK-pos, CD45-neg, and CD69-high. We used a conservative gating strategy when enumerating epithelial CD69 positivity in breast tumors to reduce possible false positives. When generating pseudocolored composite images, we assigned markers the following colors: H3NUCA (blue), CD3 (cyan), CD45 (magenta), CD69 (yellow), PANCK (red), VIM (green). Epithelial CD69 is visually portrayed as overlapping yellow and red colors.

### Cell line studies

Computational evaluation of leukocyte surface receptor expression in human breast cancer cell lines leveraged public RNA-seq data downloaded from the DepMap project (https://depmap.org/portal) (37). Most cell lines experimentally used in this study were obtained from the American Type Culture Collection. Exceptions include JIMT-1 cells, which were obtained from the DSMZ-German Collection and the PyMT-chOVA cell line originated in the Max Krummel Lab and was acquired from the Zena Werb Lab at UCSF. All cell lines were verified to be free of mycoplasma adjacent to passage numbers used in this study and this information is summarized in Table S3. The following cell lines were maintained in DMEM + 10% FBS: 4T1, CAMA-1, JIMT-1, MCF-7, MDA-MB-231, MDA-MB-468, PyMT-chOVA, and SK-BR-3. These lines were maintained in RPMI 1640 + 10% FBS: BT-474, HCC 1806, T-47D, and ZR-75-1. Cell culture medium source: DMEM (Fisher, cat #11-995-073), RPMI (Fisher, cat #11-875-093), FBS (ThermoFisher Scientific, cat. #10438026). Each cell line was propagated under subconfluent conditions and dissociated with trypsin-EDTA (0.25%) (Fisher, cat. #25-200-114) for all assays.

Drug treatments on cell lines involved paclitaxel (Sigma-Aldrich, cat. #T7402) or doxorubicin (Selleck Chemicals, cat. #S1208) solubilized in DMSO. Human breast cancer cell lines were treated with 80 nM paclitaxel or 100 nM doxorubicin, and murine mammary cancer cell lines were subjected to 200 nM paclitaxel. These concentrations aimed to achieve a half-maximal inhibitory concentration (IC50) on cell growth. All treatments were initiated 24 hours after low-density cell plating and persisted for 72 hours. In some instances, MDA-MB-231 cells were treated with differing concentrations of paclitaxel or doxorubicin to evaluate CD45 versus CD69 along a time-course. Irradiation (8 Gy dose via CIX2 Cabinet X-Ray Irradiator) and serum-free medium treatments followed this same paradigm, which began also 24 hours after plating. All cell line morphology photographs were acquired on a Nikon Eclipse Ts2R microscope in either untreated conditions or at treatment endpoint.

Cell growth was assessed via soft-agar and crystal violet assays. Evaluating colony formation *in vitro* involved plating MDA-MB-231 cells at low density and within 0.3% agar onto a 0.5% layer of agar. Half-medium exchanges were performed every 5 days for one month, after which colonies were stained with 0.005% crystal violet. Images were acquired on a dissection microscope and binarized in Fiji (ImageJ), after which colony size per well was calculated as pixel area. For general cell proliferation assays, cell growth was quantified in a 96-well vessel after crystal violet staining (0.1%), where 50 cells per well represented a sparse plating density and 500 cells per well was standard.

### Flow cytometric assessment of immune mimicry

All cell lines were dissociated with trypsin-EDTA (0.25%) (Fisher, cat. #25-200-114) for flow cytometry. Flow cytometric staining was done as monostains or all stains where indicated, and data were acquired on a 5-laser Cytek Aurora Spectral Flow Cytometer on cells fixed with BD Cytofix (BD, cat. #554655). Antibody monostains were incubated in the presence of 5% fetal bovine serum and all stains were incubated in Brilliant Stain Buffer (BD Biosciences, cat. #563794). The antibodies used in this study are listed in Table S4 (human) and Table S5 (murine). KI67 staining was performed on cells following fixation/permeabilization (ThermoFisher Scientific, cat. #88-8824-00). We included mouse-CD45-PerCP when conducting flow cytometry on digested organs containing mCherry-pos MDA-MB-231, and all flow cytometric staining on dissociated mouse tissues followed FC Block (BD Biosciences, cat. #553142). Flow cytometry done on CD69 CRISPR-a MDA-MB-231 cells necessitated changing marker fluorophores to accommodate high eGFP levels in the cell line; we used Zombie Violet and CD69-APC and ran these samples on a BD LSRFortessa. Flow cytometric data were analyzed in FCS Express, where we applied upstream gates to exclude cellular doublets, dead cells, and high autofluorescence. The flow cytometry marker correlation plot was made with the corrplot R package (RRID:SCR_023081) and tSNE plots were generated with the Spectre R package (38).

### qRT-PCR assessment of immune mimicry

The qRT-PCR primers used in this study are shown in Table S6. RNA extractions were performed with an RNeasy kit (Qiagen, cat. #74104) and cDNA was synthesized using SuperScript VILO Master Mix (ThermoFisher Scientific, cat. #11-755-050) or iScript (Bio-Rad, cat. #1708891). Quantitative PCR was conducted on an Applied Biosciences ViiA 7 with PowerUp SYBR Green (ThermoFisher Scientific, cat. #A25741) or on a BioRad Opus with iTaq Universal SYBR Green (Bio-Rad, cat. #1725121). Gene expression in cell lines was normalized to either ACTB or GAPDH. CD69 expression after CRISPR/Cas9 editing was evaluated with primers binding upstream of the sgRNA target sites. All primer oligos were purchased from Integrated DNA Technologies.

### CRISPR editing and shRNA

All sgRNA sequences used to overexpress or disrupt CD69 are found in Table S7 and plasmids are listed in Table S8. All CD69 sgRNA oligos were identified in the CHOPCHOP database (https://chopchop.cbu.uib.no/) and obtained from Integrated DNA Technologies. CD69 sgRNAs were cloned using the annealed oligo method into their requisite vector backbones. Endogenously overexpressing CD69 employed the CRISPR Synergistic Activation Mediator system. MDA-MB-231 cells were stably transduced with the following plasmids for CRISPR-activation experiments: pHAGE TRE dCas9-VP64, (AddGene plasmid #50916), lenti sgRNA(MS2)_Puro backbone (AddGene plasmid #73795), LentiMPH v2 (AddGene plasmid #89308), and pLV-eGFP (AddGene plasmid #36083) (39–42). MDA-MB-231 cells were treated with 1μg/ml doxycycline to induce CD69 during CRISPR-activation experiments. MDA-MB-231 cells were stably transduced with lentiCRISPR v2 (AddGene plasmid #52961) for CD69 CRISPR/Cas9 disruption studies (41). To deplete CD45, MDA-MB-231 cells were stably transduced with a GIPZ shRNA viral particle kit (Revvity Discovery, shRNA-1 cat. #VGH5518-200199261, shRNA-2 cat. #VGH5518-200261641) and compared to cells transduced with the manufacturer’s non-silencing control.

### Magnetic cell sorting

CD69-high neoplastic cells were sorted from MDA-MB-231 and 4T1 cultures using an EasySep PE Positive Selection Kit II (STEMCELL Technologies, cat. #17684). We used human-CD69-PE (BioLegend cat. #310905) and mouse-CD69-PE (BioLegend cat. #104512) antibodies for these experiments. Each sorting replicate was conducted on cultured cells grown under standard conditions for 72h hours to a final confluency of ∼75%. Sorting 200 million cells per replicate, on average, yielded around 50,000 CD69-high cells and thereby limited downstream applications.

### Orthotopic *in vivo* tumor studies

Orthotopic tumor growth was evaluated in all cell lines implanted into the 4^th^ left mammary gland of female mice between 6-8 weeks of age. Female NSG mice (Jackson Laboratory, cat. #005557, RRID:IMSR_JAX:005557) received 1 million MDA-MB-231 cells suspended in DMEM containing 50% Matrigel GFR Basement Membrane Mix (Corning, cat. #354230). Mice used for CD69 CRISPR-a experiments were fed a 200 mg/kg doxycycline diet (Bio-Serv cat. #S3888) that was initiated 8 days after MDA-MB-231 implantation. Metastases were analyzed in H&E lung sections using QuPath to quantify the size of individual lesions and the number of cells within them. Orthotopic experiments with 4T1 cells sorted on CD69 involved implanting 1,000 cells orthotopically into BALB/c mice (Charles River, cat. #028, RRID:IMSR_CRL:028) in PBS. In some instances, an mCherry fluorescent tag (AddGene plasmid #36084) was stably transduced into 4T1 and MDA-MB-231 cells to facilitate their retrieval from tissue *in vivo*. In these studies, we implanted 200,000 4T1 cells in PBS into BALB/c mice or 1 million MDA-MB-231 cells in Matrigel GFR into NSG mice. For orthotopic reinjection experiments with dissociated MMTV-PyMT-chOVA-CD69^KO/+^ or -CD69^KO/KO^ tumors, we first selected tumors from each background that: developed in the same mammary gland, grew at similar rates, and reached a similar size at endpoint. We found three pairs of tumors that we reinjected into CD69^KO/+^ mice lacking the PyMT transgene. When required, mammary tumors were digested in DMEM containing 2 mg/ml Collagenase A (Roche cat. #11088793001) and 50 U/ml DNase I (Roche cat. #10104159001) with agitation for 30 min at 37°C. Experiments attained endpoint when tumors reached 2 cm in diameter, if they became ulcerated, or if mice exhibited outward signs of suffering.

### Genetically engineered *in vivo* tumor studies

Spontaneous mammary tumor growth was assessed in MMTV-PyMT-chOVA-CD69^KO/KO^ mice that were generated by crossing MMTV-PyMT-chOVA mice (43) with CD69^KO/KO^ mice (44,45). Knockout of the CD69 gene was routinely confirmed through genotyping, and also demonstrated on mouse peripheral blood by flow cytometry in a subset of animals following anti-CD3 (BioLegend, cat. #100340, RRID:AB_11149115) stimulation for 4 hours. Mice were monitored for mammary tumors starting at 100 days, and at 3/4-day intervals thereafter until endpoint. We recorded tumor incidence and size data that were compared between MMTV-PyMT-chOVA-CD69^KO/+^ and -CD69^KO/KO^ mice. Tumor growth was evaluated between these groups and doubling time was calculated from an exponential regression model, tumor slope from a linear regression model, and both were subjected to outlier removal in GraphPad. Tumor burden per mouse was calculated by summing all tumors, and we subjected these data to nonlinear regression (Gompertz growth) to calculate when tumor burden crossed the 1000 mm^3^ threshold. For all MMTV-PyMT experiments, mice were euthanized when tumors reached 2 cm in diameter, if the tumors became ulcerated, or if mice showed visible signs of suffering.

### Statistical analyses

Statistical analyses were carried out with GraphPad Prism v10. We considered the distribution of data points when conducting statistics and the tests used in this study include: analysis of variance, linear regression, nonlinear regression, Pearson correlation, Spearman correlation, Kruskal-Wallis test, Mann-Whitney test, Wilcoxon signed-rank test, Student’s t-test, and Mantel-Cox log-rank test. All experimental data points are shown as biological replicates.

### Data availability

Human breast tumor atlas scRNA-seq datasets (23,24) are publicly available from the Gene Expression Omnibus (Pal *et al.*, GSE161529; Wu *et al.*, GSE176078) as are reduction mammoplasty atlas data (23,24) (Pal *et al.*, GSE161529; Kumar *et al.*, GSE235326). PANNTHR trial scRNA-seq data were deposited in the NCBI Sequence Read Archive under BioProject accession number PRJNA1309720. The PBMC dataset was downloaded through the website accompanying its publication (31). The R code involved in the scRNA-seq analysis is located at https://github.com/EBBerens/Neoplastic-immune-mimicry-potentiates-breast-tumor-progression.

## Results

### Identification of neoplastic immune mimicry in human breast tumors and reduction mammoplasties

The premise of our study was that neoplastic cells engaging in plasticity could be computationally identified in breast cancer. We hypothesized that these neoplastic cell states, if detected, likely contribute to disease progression and would therefore be present in multiple tumors. Our ensuing Seurat-based (28) scRNA-seq analysis leveraged publicly available human breast tumor datasets (23,24). We queried whether stem-like neoplastic cells cluster with, or separately from, well-differentiated epithelium and thus developed a computational approach to detect them irrespective of their cluster location (**Fig. 1A**). This modified approach isolated neoplastic mammary epithelium by first removing multiplets (29), thereafter selecting cells expressing cytokeratin biomarker genes (e.g. KRT14, KRT18, and KRT19) along with inferred copy number variations (CNVs). We carried out scRNA-seq preprocessing on 70 human breast tumor samples derived from three datasets: the Pal *et al.* atlas of 34 therapy-naïve tumors (23) (**fig. S1A-G)**, the Wu et al. atlas of 26 tumors (*24*) (**fig. S2A-G)**, and 10 tumor samples from the PANNTHR clinical trial at the Oregon Health & Science University (**fig. S3A-G**). After scRNA-seq preprocessing, our analysis considered 40 breast tumors with hormone receptor-positive disease, 11 with HER2-amplified disease, and 18 with triple-negative disease; 59 of these tumors were therapy-naïve at the time of sample collection and 10 had seen therapy. An atlas of 124 reduction mammoplasties (27) was also analogously subjected to scRNA-seq preprocessing but without CNV detection (**fig.S4A-F**) after which merged datasets were generated for downstream analyses.

**Figure 1.**
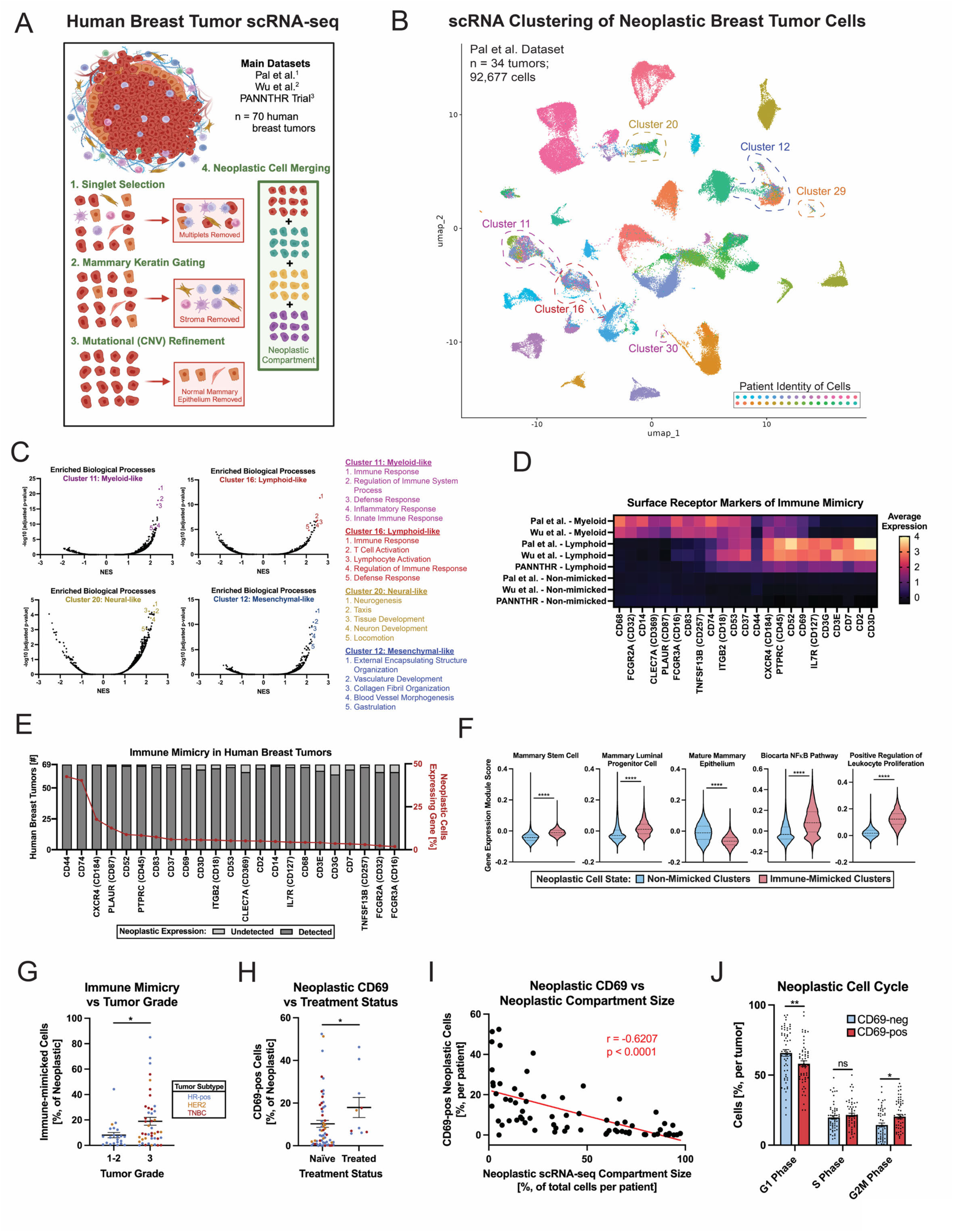
Neoplastic immune mimicry is identified in independent human breast tumor scRNA-seq cohorts. (**A**) scRNA-seq workflow to collect neoplastic cells based on mammary keratin gene expression. (**B**) Example of scRNA-seq clustering from the Pal *et al*. dataset where cells are colored according to their patient identity. Indicated clusters depict neoplastic cell states shared across patients that (**C**) express myeloid-like, lymphoid-like, neural-like, and mesenchymal-like gene signatures. (**D**) Immune mimicry is signified by neoplastic cells expressing canonical leukocyte surface receptor genes in a concordant manner across distinct scRNA-seq datasets. (**E**) The vast majority of breast tumors analyzed have neoplastic cells expressing these immune mimicry markers as a minority subpopulation. (**F**) Neoplastic cells found within immune-like scRNA-seq clusters are enriched for dedifferentiated, NFκB pathway, and proliferative gene signatures. (**G**) Neoplastic immune mimicry is elevated in high-grade breast tumors. Breast tumors grade 1-2, n = 22; grade 3, n = 44; tumors with both pre- and post-treatment data depicted once. (**H**) Neoplastic cells expressing the CD69 early activation marker are increased in tumors subjected to therapy before surgical resection, (**I**) in scRNA-seq samples harboring a smaller epithelial compartment, and (**J**) they are more often classified as mitotic. In (H), therapy-naïve tumors, n = 59; treated tumors, n = 10. In (I), 69 breast tumors shown. In (J), 58 tumors with at least 10 CD69-pos neoplastic cells depicted. * p < 0.05; ** p < 0.01; **** p < 0.0001. Wilcoxon signed-rank test (F), Mann-Whitney test [(G) and (H)], Pearson correlation (I) or two-way ANOVA with Šidák correction (J). Data are means ± SEM.

Evaluating neoplastic epithelia in 34 therapy-naïve breast tumors from the Pal *et al.* dataset (23) revealed distinct neoplastic scRNA-seq clusters mostly formed by cells from individual patients, presumably driven by interpatient variability (**Fig. 1B; fig. S5A-B**). We used Shannon-Weaver Diversity Index scoring to identify clusters comprised of neoplastic cells amongst different patients, reasoning such clusters could represent shared cell states. Six shared clusters contained elevated Shannon-Weaver Diversity Index scores along with evident enrichment of unexpected non-epithelial biological processes typically attributed to immune (myeloid or lymphoid), neural, mesenchymal, and endothelial cells (**Fig. 1C**; **fig. S5C-E**). These non-epithelial biological processes were largely corroborated within shared neoplastic clusters in the other human breast tumor scRNA-seq datasets (**fig. S6A-E; fig. S7A-E)**. We were particularly intrigued by the possibility that neoplastic epithelia expressed genes ascribed to leukocytes and investigated this program further. In doing so, we focused specifically on genes encoding cell surface receptors due to their potential clinical applicability as future biomarkers or therapeutic targets.

Upregulated within these shared neoplastic clusters were 23 surface receptor genes significantly associated with “myeloid-like” and/or “lymphoid-like” neoplastic cell phenotypes across the breast tumor scRNA-seq datasets (**Fig. 1D; fig. S5F; fig. S6F; fig. S7F**). Several genes notably encode proteins routinely used to distinguish leukocyte subsets: e.g. CD3, CD14, CD18, CD45, CD68, CD69, CD74, and CD83. In considering all the breast tumors analyzed, we learned that neoplastic cells expressing these immune biomarkers were detected in most samples yet only as minority (<5%) subpopulations on average (**Fig. 1E**). Because CD45 (PTPRC) was considered a pan-leukocyte receptor and canonically not expected in neoplastic cells, we compared inferred CNVs in CD45-pos vs CD45-neg cells as another layer of neoplastic validation. CD45-pos neoplastic cells indeed had gene- and region-level inferred CNVs akin to CD45-neg neoplastic cells (**fig. S8A-D**) and on a patient-by-patient basis (**fig. S8E-F**), supporting our hypothesis that immune mimicry was neoplastic in origin. We next evaluated neoplastic epithelial cells expressing the “myeloid-like” and “lymphoid-like” genes from all datasets and therein broadly refer to them as “immune-mimicked” going forward.

Immune-mimicked cells in human breast tumors exhibited significantly upregulated signatures indicative of mammary stem and progenitor cells (46) including the NFκB pathway, and proliferating leukocytes when compared to non-mimicked neoplastic epithelia (**Fig. 1F**; **fig. S5G; fig. S6G; fig. S7G**). We considered whether the NFκB pathway genes might represent a conduit enabling neoplastic cells to activate immune surface receptor expression, given its dual roles in regulating immune responses as well as malignant cell behaviors (47). These observations intersected with increased neoplastic immune mimicry in high-grade breast tumors (**Fig. 1G**). Importantly, we found evidence of immune mimicry in presumably non-neoplastic epithelial cells obtained from reduction mammoplasties (27) indicating that this state is not restricted to malignant epithelium (**fig. S9A-G**), however and of interest, immune mimicry was significantly amplified in mammoplasties collected from patients with a high body mass index (**fig. S9H**) which is a risk factor for developing aggressive breast cancer (48). These data thereby denoted the presence of immune mimicry in both homeostatic and diseased breast tissue.

Neoplastic cells often coexpressed multiple leukocyte surface receptors (**fig. S10A-B**) coincident with the variable upregulation of known breast tumor-initiation biomarkers (**fig. S10C**). Immune mimicry was more pronounced in breast tumors versus reduction mammoplasties (**fig. S10D**) with the greatest level of immune mimicry found in triple-negative breast cancer (TNBC) (**fig. S10E**). Comparing RNA expression for immune mimicry genes in neoplastic cells versus epithelia from reduction mammoplasties revealed an association between CD69 and malignant epithelia (**fig. S10F-G**). CD69 is a canonical leukocyte activation marker highly expressed on T cells where it enhances their retention in tissues and other functions including proliferation (25,26). We posited a parallel function for epithelial CD69 in breast cancer and selected it for follow-up studies, reasoning that establishing CD69 in neoplastic cells could have utility in future clinical diagnostics.

Because CD69 is typically activated in leukocytes during inflammation and tissue stress (25), we accordingly hypothesized that epithelial CD69 might arise in tumors subjected to cytotoxic therapy. We thus compared breast tumors receiving cytotoxic therapy prior to resection versus those that were therapy-naïve where significantly elevated neoplastic CD69 expression followed therapy (**Fig. 1H; fig. S10H**), and tumors containing fewer neoplastic cells (**Fig. 1I**), implying enrichment in cells responding poorly to treatment or subsisting thereafter. CD69-positive neoplastic cells in breast tumors were additionally often classified as mitotic (**Fig. 1J**) based on cell cycle phase scoring performed in Seurat (28). These observations engendered the possibility that neoplastic epithelia express CD69 amid cell stress to facilitate tumor progression.

To further address whether immune mimicry and neoplastic CD69 represented an artifact, we next verified that immune-mimicked cells reflected non-mimicked neoplastic epithelia concerning their RNA features, RNA counts, mitochondrial reads, (**fig. S11A**) and doublet probability score (**fig. S11B**). These results implied that immune mimicry was unlikely to ensue from dying or aggregated cells. Neoplastic cells expressing canonical leukocyte surface receptor genes were found outside and within immune-mimicked scRNA-seq clusters (**fig. S11C**), indicating the phenotype resided along a spectrum. Hybrid cells within the immune-mimicked clusters ostensibly upregulated leukocyte activation processes versus non-mimicked neoplastic cells (**fig. S11D**) while retaining aspects of epithelial differentiation versus leukocytes in general (**fig. S11E**). Lastly, we interrogated a ∼2 million human peripheral blood mononuclear cell (PBMC) public dataset (31) in search of epithelial cells and discovered a small fraction expressing KRT18 though otherwise devoid of KRT14 and KRT19 (**fig. S11F-I**). Thus, the mammary cytokeratin profile of immune-mimicked cells was more closely related to non-mimicked neoplastic epithelia than PBMCs (**fig. S11I**). We next sought to validate our aforementioned scRNA-seq findings in additional settings, starting with the histopathological assessment of immune mimicry and epithelial CD69 in human breast tumor specimens.

### Histopathologic confirmation of epithelial CD69 in human breast tumors

The Human Protein Atlas (35) contained histopathological images for immune mimicry protein staining across human breast tumors. Exploration of these public data revealed convincing instances of leukocyte surface receptor staining in the neoplastic compartment for CD14 and CD74 that coincided with the literature (15). Our search also uncovered samples harboring neoplastic CD83, as well as limited cases of prospective weak detection in neoplastic cells for CD3 and CD68. Absent from the Human Protein Atlas were any examples of neoplastic CD45 and CD69, thus we pursued their validation in-house and evaluated epithelial CD45 and CD69 in 58 histopathological human breast tumors via multiplexed immunohistochemistry (mIHC).

Unbiased quantification of neoplastic CD45 in breast tumors was challenging due to leukocyte infiltrates often abutting neoplastic cells. Nevertheless, visual inspection indicated possible areas of epithelial-mesenchymal transition (12) within high-grade breast tumors where CD45-positive (yellow), CD68-positive (cyan), and vimentin-positive (green) cells bordered pan-cytokeratin (red) epithelium (**fig. S12A-C; fig. S13A**). In some regions positive cells for both CD45 and pan-cytokeratin were discerned (**fig. S12B**). The CD45-positive cells in question exhibited morphology, heterotypic junctions, and a nuclear size that seemed reminiscent of neighboring epithelium, however we deemed these data too ambiguous to render a clear verdict.

Our subsequent analysis of neoplastic CD69 revealed clear minor subpopulations positive for both CD69 (yellow) and pan-cytokeratin (red) in several breast tumors (**Fig. 2A-B**). Here, neoplastic CD69 expression patterns ranged from heterogeneous and diffuse to concentrated depending on the histopathological region of cytokeratin-positive cells (**Fig. 2B**). To limit false-positives, we only considered epithelium with strong CD69 staining which led to positivity identified in over half of the tumor cohort (**Fig. 2C**) with no statistical tropism for disease subtype (**fig. S14A-D**). There was a trend toward increased neoplastic CD69 in tumors treated with cytotoxic therapy prior to resection (**fig. S14E**) where CD69-positive epithelial cells trended toward greater positivity for Ki67 more frequently than CD69-neg cells (**fig. S14F-H**). Given the histopathological confirmation of neoplastic CD69 expression and possible evidence for other immune mimicry markers, we hypothesized that breast cancer cell lines would likewise express canonical leukocyte surface receptor genes and proteins.

**Figure 2.**
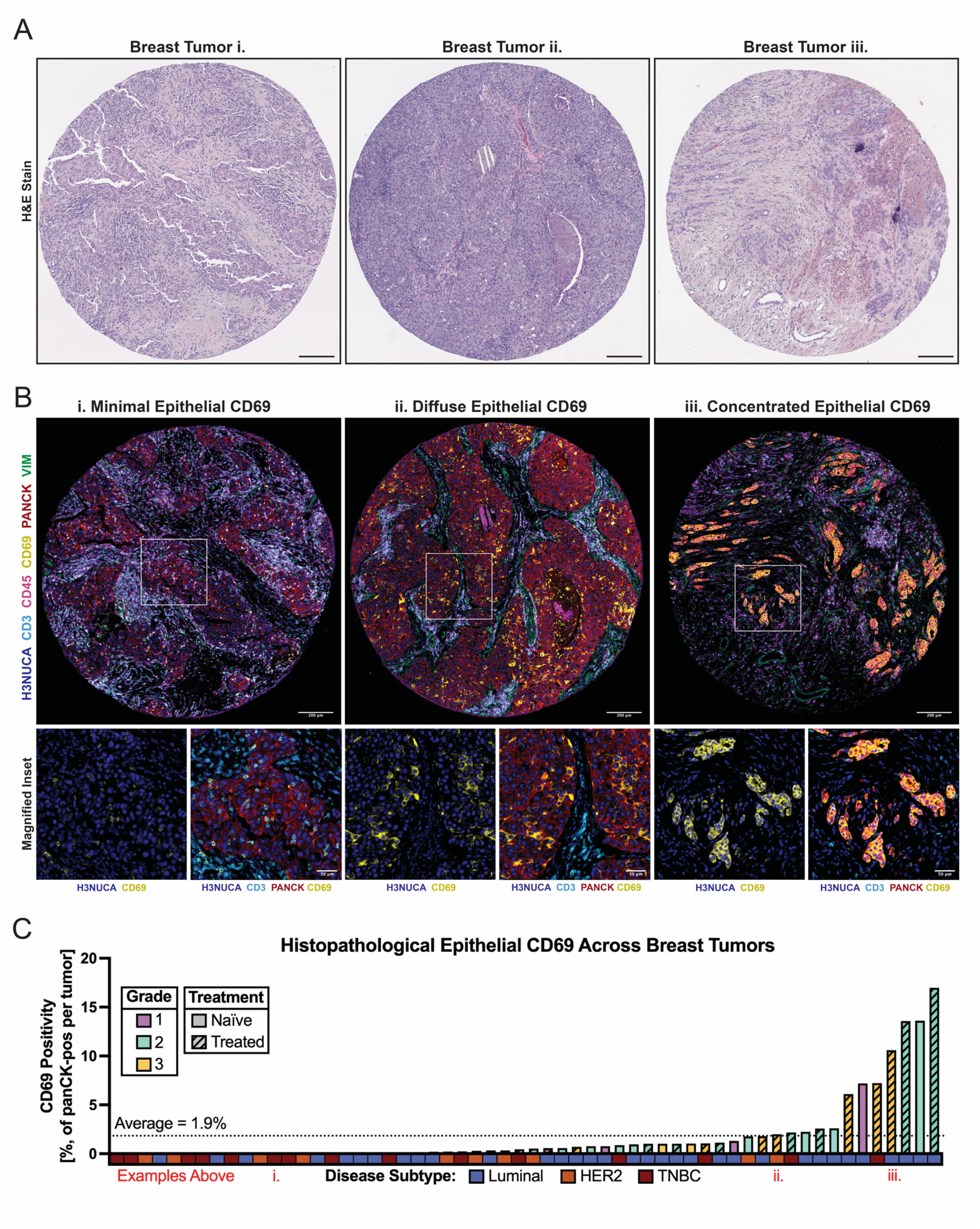
The immune mimicry CD69 biomarker is associated with aggressive breast cancer in histopathological samples. (**A**) Hematoxylin and Eosin (H&E) staining of human breast tumors. These same tissues were subsequently subjected to multiplexed immunohistochemistry to detect epithelial CD69. Scale bar, 200 μm. (**B**) Multiplexed immunohistochemistry examples of neoplastic CD69 or the lack thereof in primary human breast tumors. Neoplastic CD69 is denoted by overlapping PANCK (red) and CD69 (yellow). The first example portrays a lack of neoplastic CD69 amid immune cell (CD3-pos, cyan) expression. The second and third examples show neoplastic CD69 in diffuse and concentrated staining patterns, respectively. Scale bar, 200 μm. The insets provide magnified views of CD69 (left) versus CD3 and pan-cytokeratin (right). Inset scale bar, 50 μm. (**C**) Numerous human breast tumors have detectable levels of neoplastic CD69. Plot depicts the percent of epithelial cells strongly positive for CD69 for distinct tumors along with clinical parameters. Epithelial CD69 is summarized from a conservative gating approach applied to multiple regions in fifty-eight tumors.

### Neoplastic immune mimicry in breast and mammary cancer cell lines

Our cell line analysis began by probing public bulk RNA-seq data from the Cancer Dependency Map (DepMap) (37) for evidence of immune mimicry surface marker gene expression. We initially focused our efforts on 57 breast cancer cell lines available through DepMap that represented the major disease subtypes, noting prevalent leukocyte marker RNA expression at baseline (**Fig. 3A**). In these data, immune mimicry genes were significantly elevated in basal breast cancer cell lines compared to luminal or HER2-amplified subtypes (**Fig. 3B**) and significantly correlated with dedifferentiation and NFκB pathway activation signatures (**Fig. 3C-E**). We next experimentally validated a subset of the immune mimicry receptors as protein monostains in 10 human breast cancer cell lines using flow cytometry. Consistent with scRNA-seq analyses, these efforts unveiled minority subpopulations of viable cells expressing canonical immune surface receptors (e.g. CD3, CD14, CD18, CD45, CD69, CD74, CD83) that varied across biological replicates (**Fig. 3F-G**). Basal subtype cell lines exhibited greater average leukocyte receptor expression (**Fig. 3H; fig. S15A-B**) in agreement with DepMap RNA data (**Fig. 3A**). By flow cytometry, we discovered CD45-positive neoplastic cells were significantly smaller in size than CD45-negative cells (**Fig. 3I**), and additional co-staining revealed that CD45 often correlated with expression of other immune mimicry markers including CD52, CD69, CD74, and CD83 (**Fig. 3J**). Because CD45-positive cells were not enriched for classic tumor-initiation markers, e.g., CD44, CD133, KIT, or THY-1, we concluded that immune-mimicked cells either lacked stem-like features *in vitro* or they were dedifferentiated in a different manner (**fig. S15C**). Nevertheless, these immune-mimicked cells represented novel subpopulations worthy of additional study.

**Figure 3.**
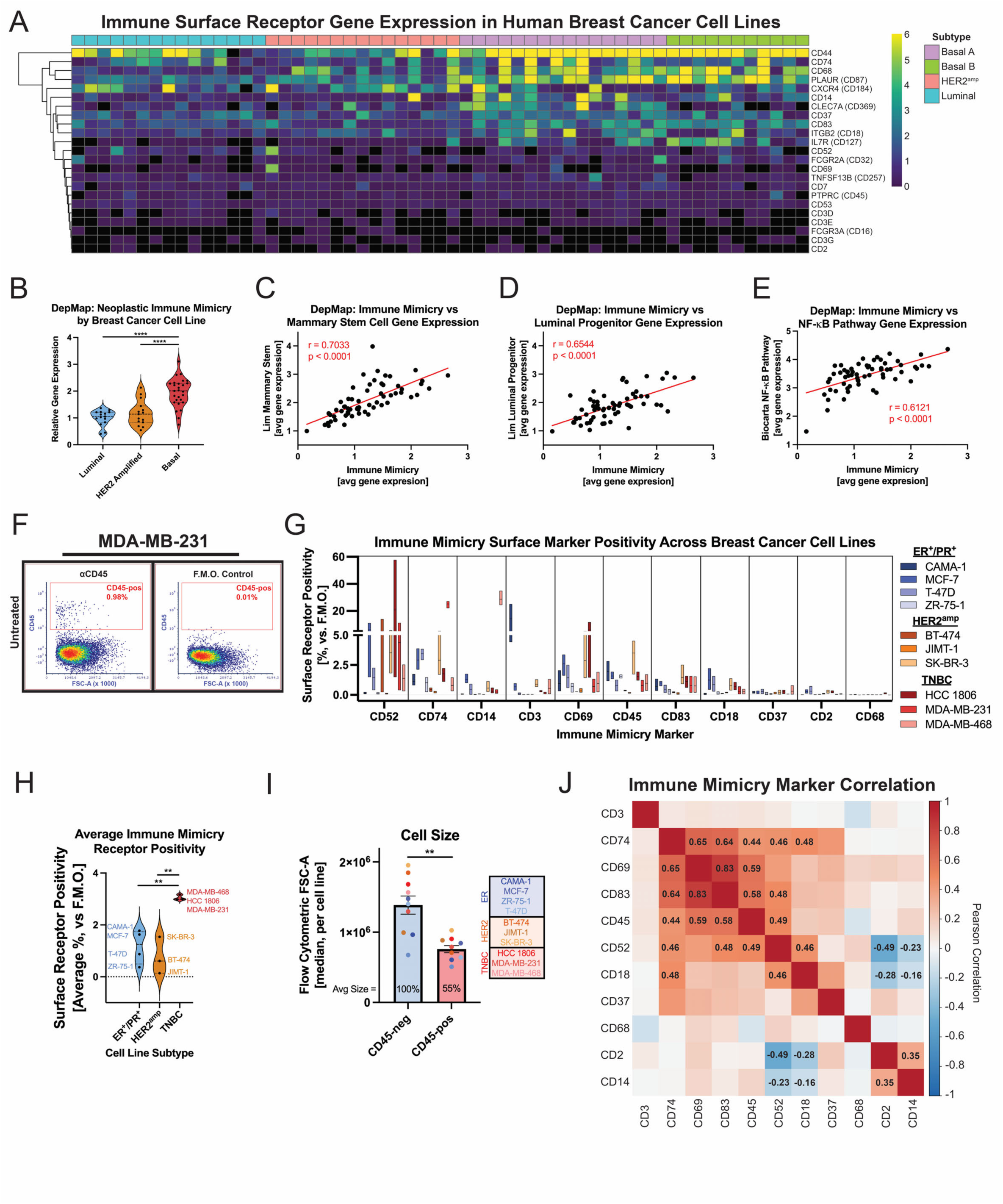
Breast cancer cell lines express leukocyte surface receptor RNA and protein. (**A**) Interrogation of immune mimicry gene expression in public bulk RNA-seq DepMap data encompassing fifty-seven breast cancer cell lines. (**B**) Neoplastic immune mimicry RNA gene expression is highest in breast cancer cell lines of the basal phenotype. Breast cancer cell lines expressing greater levels of leukocyte surface receptor genes positively correlate with (**C**) mammary stem, (**D**) mammary luminal progenitor, and (**E**) NFκB pathway gene signatures. (**F**) Example of flow cytometric CD45 protein staining in MDA-MB-231 cells. Negative (fluorescence minus one) control shown for comparison. (**G**) Demonstration of canonical leukocyte surface receptor protein positivity in ten human breast cancer cell lines via flow cytometry. Data are monostains performed in biological triplicate. (**H**) Triple-negative breast cancer cell lines exhibit greater levels of leukocyte surface receptor protein. (**I**) Immune-mimicked CD45-pos cells are significantly smaller than CD45-neg cells. (**J**) Immune mimicry marker correlations across the ten cell lines subjected to flow cytometry analysis. Coefficients for significant correlations are indicated. ** p < 0.01; **** p < 0.0001. One-way ANOVA with Tukey correction [(B) and (H)], Pearson correlation [(C), (D), (E) and (J)], or Wilcoxon signed-rank test (I). Data are means ± SEM.

### Neoplastic CD69 denotes a cell state capable of proliferating under stress conditions

We hypothesized that leukocyte receptor positivity would increase in cell lines subjected to stress derived from cytotoxic therapy or serum starvation, based on observations for neoplastic CD69 in scRNA-seq and mIHC data (**Fig. 1H; fig. S14E**). Indeed, CD69 cell positivity increased in aggressive/metastatic MDA-MB-231 and HCC 1806 cell lines following paclitaxel despite all models possessing a similar baseline (**Fig. 4A; fig. S16A-D**). A time-course experiment revealed that surface CD45 and CD69 gradually increased on MDA-MB-231 cells during 72 hours of paclitaxel exposure (**fig. S16E**). Using MDA-MB-231, we quantified leukocyte surface receptor expression in various additional cytotoxic stress conditions and uncovered consistently elevated positivity for CD45, CD69, and CD83 (**Fig. 4B**). Several proteins displayed an affinity for the manner of cell stress, exemplified by MDA-MB-231 CD3 enrichment in serum-free medium and in response to doxorubicin, but not in response to paclitaxel (**Fig. 4B**). To further demonstrate that immune mimicry was not artifactual, we successfully depleted CD45 using short hairpin RNA (shRNA) in MDA-MB-231 cells amid doxorubicin (**fig. S17A-D**) and also found that higher drug concentrations led to loss of CD45, indicating the phenotype required viable cells (**fig. S17E**).

**Figure 4.**
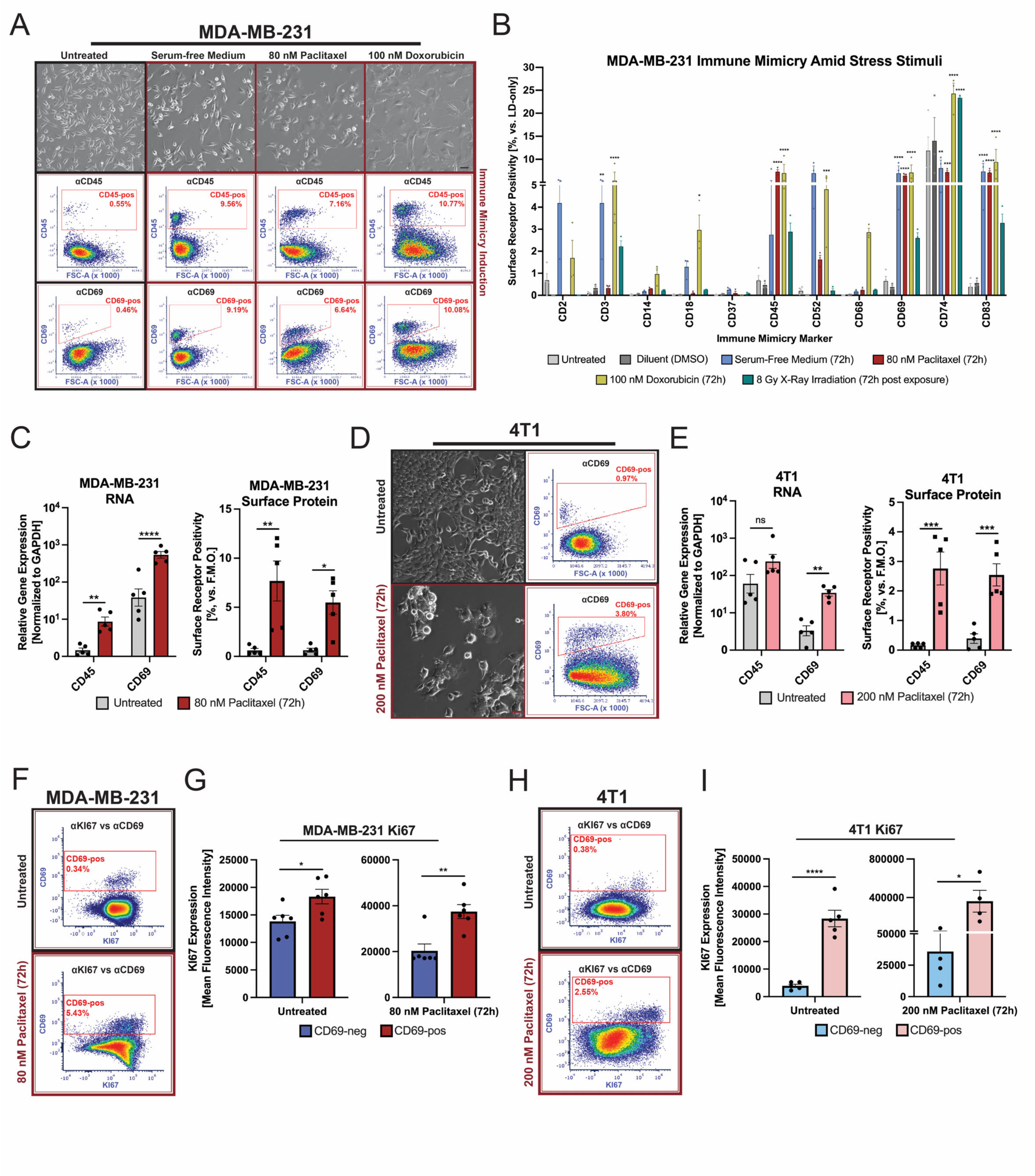
Cytotoxic therapy further induces neoplastic immune mimicry and CD69. (**A**) Cell stress stimuli robustly expand the subpopulation of cells expressing CD45 and CD69 in MDA-MB-231. Scale bar, 50 μm. (**B**) Example of immune surface receptor positivity for MDA-MB-231 cells across various cytotoxic conditions. Data represent flow cytometric analysis of all stains performed in biological triplicate. (**C**) CD45 and CD69 are induced at the RNA and protein level in MDA-MB-231 in response to paclitaxel. Data reflect five biological replicates with surface protein measured as flow cytometry monostains. (**D**) Flow cytometric example of paclitaxel treatment expanding the CD69-pos subpopulation in murine mammary 4T1 cells. Scale bar, 25 μm. (**E**) CD45 and CD69 are transcriptionally elevated in 4T1 cells following paclitaxel treatment. Data reflect five biological replicates with surface protein measured as flow cytometric monostains. (**F**-**G**) CD69-pos MDA-MB-231 cells express higher levels of Ki67 in untreated and paclitaxel-treated culture conditions. Data reflect six biological replicates. (**H**-**I**) CD69-pos 4T1 cells express higher levels of Ki67 in untreated and paclitaxel-treated culture conditions. Data reflect five or four biological replicates, respectively. * p < 0.05; ** p < 0.01; *** p <0.001; **** p < 0.0001. Two-way ANOVA with Dunnett correction (B) or Šidák correction [(C) and (E)], or Student’s t-test [(G) and (I)] Data are means ± SEM.

Paclitaxel-induced CD45 and CD69 in MDA-MB-231 occurred at both RNA and protein levels implying transcriptional activation or selective survival compared to baseline (**Fig. 4C; fig. S18A-B**). This phenotype contrasted less aggressive MCF-7 cells in which immune mimicry was not impacted by paclitaxel (**fig. S18C-D**). Akin to MDA-MB-231, aggressive murine mammary 4T1 cells also had expression of immune surface receptors following paclitaxel treatment (**fig. S18E-F**), with both CD69 RNA and protein being enhanced (**Fig. 4D**). Moreover, immune-mimicked cells expressing CD69 had higher levels of Ki67 in both untreated and paclitaxel-treated conditions for MDA-MB-231 (**Fig. 4F-G**) and 4T1 (**Fig. 4H-I**). Together, these data support the notion that neoplastic CD69 distinguishes a cell state capable of proliferating under stress conditions.

Since harsh conditions are naturally elicited during tumor metastasis, we asked whether neoplastic CD45 and CD69 were increased on 4T1 and MDA-MB-231 in this context. Each cell line was transduced with an mCherry fluorescent reporter to facilitate *in vivo* detection and then subjected to a tumor growth and metastasis assay in mice, followed by tissue collection and flow cytometry analyses (**fig. S19A**). At study endpoint, 4T1 mCherry cells were prospectively identified in single-cell suspensions isolated from tumor, blood, and lung tissue via co-expression of mCherry with the CD24 and EPCAM epithelial markers (**fig. S19B-C**). Disseminated 4T1 cells in the blood and lungs of BALB/c mice significantly upregulated CD45 expression, while CD69 was tropic for cells isolated from metastatic lungs (**fig. S19D**). Conducting a parallel study with MDA-MB-231 in immunocompromised NSG mice involved prospective retrieval using mCherry and CD44 (**fig. S19E-F**) where we observed MDA-MB-231 metastases in numerous organ sites that expressed human CD45 and CD69 (**fig. S19G**). Strikingly, in metastatic lungs, MDA-MB-231 CD69 declined as metastatic burden increased (**fig. S19H**) which raised the possibility that neoplastic cells might coopt CD69 for the early stages of metastatic outgrowth, analogous to how leukocytes utilize CD69 for cell activation (25,26).

### Experimental studies linking neoplastic CD69 to enhanced early tumor growth

To evaluate links between neoplastic CD69 and malignancy, we employed CRISPR-activation (CRISPR-a) to endogenously overexpress CD69 in MDA-MB-231 cells, hypothesizing this would enhance aggressive cell proliferation *in vitro* and *in vivo*. Activation of the doxycycline-induced CRISPR-a system led to prominent CD69 RNA and protein expression (**Fig. 5A-C**). MDA-MB-231 cells overexpressing CD69 formed larger colonies during low-density expansion on soft-agar indicating enhanced proliferation/survival *in vitro* (**Fig. 5D-E**). Orthotopic implantation of MDA-MB-231 cells into immunocompromised NSG mice and the subsequent activation of CD69 (**Fig. 5F**) notably accelerated early tumor growth with respect to the CRISPR-a control (**Fig. 5G-H; fig. S20A-C**), with the difference dissipating by study endpoint (**Fig. 5I-J; fig. S20D-F**). However, at endpoint, mice implanted with activated-CD69 MDA-MB-231 cells had significantly larger metastatic lung foci versus the control (**Fig. 5K-L**) despite no difference in overall number of lung lesions (**Fig. 5M**). These data implied that neoplastic CD69 was not likely impacting lung seeding, but instead the proliferative potential of early tumors and metastases.

**Figure 5.**
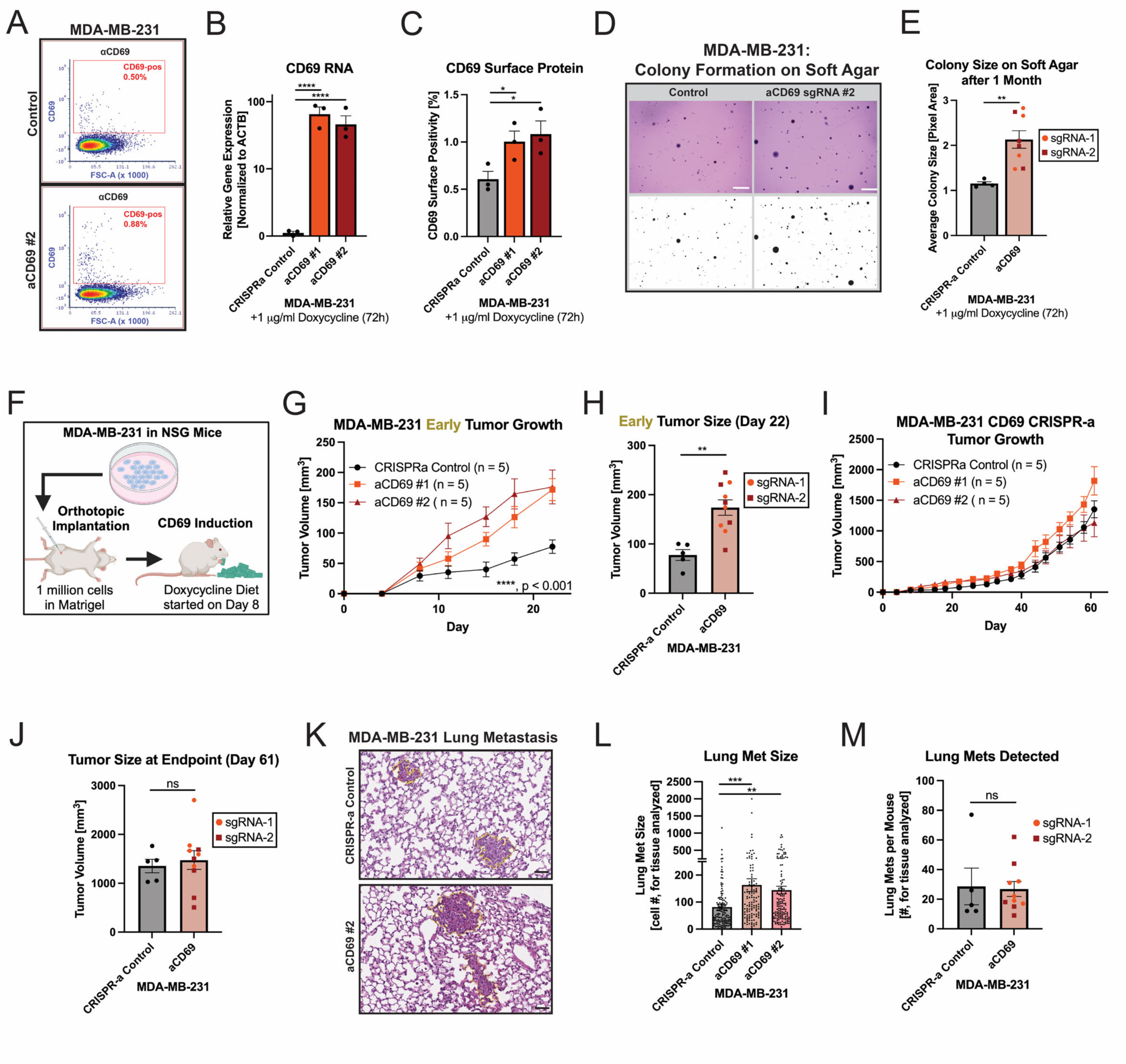
MDA-MB-231 cells overexpressing CD69 exhibit increased growth at low density *in vitro* and in mice. (**A**) CRISPR-activation of CD69 via doxycycline exposure in MDA-MB-231 increases its (**B**) RNA and (**C**) surface protein expression. Data reflect three biological replicates. (**D-E**) CD69-activated MDA-MB-231 cells form larger colonies when plated at low density on soft-agar. Data reflect four biological replicates per condition. Scale bar, 2.0 mm. (**F**) MDA-MB-231 cells were orthotopically implanted into NSG mice for a tumor growth and metastasis assay. Mice were subsequently fed a doxycycline diet on day-8 to induce CD69 in implanted cells. (**G-H**) CD69-activated MDA-MB-231 cells demonstrated enhanced early tumor growth that (**I-J**) abated by the study endpoint. Five mice per group. (**K**) At endpoint, NSG mice implanted with CD69-activated MDA-MB-231 cells had lung metastases that (**L**) were larger but (**M**) not more numerous. Individual lesions depicted in (L). Five mice per group. Scale bar, 50 μm. ns = not significant, * p < 0.05; ** p < 0.01; *** p <0.001; **** p < 0.0001. One-way ANOVA with Tukey correction [(B) and (C)] or Dunnett correction (L), Student’s t-test [(E), (H) and (J)], two-way ANOVA [(G) and (I)], or Mann-Whitney test (M). Data are means ± SEM.

We further investigated the impact of “high” versus “low” CD69 expression by utilizing magnetic bead sorting to compare the proliferative potential of CD69-high vs CD69-low subpopulations in MDA-MB-231 and 4T1 (**Fig. 6A**). Endogenous retrieval of CD69-high fractions was highly inefficient due to the small subpopulations in routine culture conditions and thus required sorting of hundreds of millions of cells per replicate. CD69-high MDA-MB-231 cells exhibited RNA enrichment for both CD69 and NFκB1 (**Fig. 6B-C**), showed enrichment trends for other NFκB pathway genes (**fig. S21A-E**), and had improved growth kinetics when plated at low density *in vitro* (**Fig. 6D**). Magnetically-sorted CD69-high 4T1 cells were similarly enriched for CD69 and NFκB1 RNA expression (**Fig. 6E-F**) along with other NFκB pathway genes (**fig. S21F-J**). We orthotopically implanted CD69-high and CD69-low 4T1 cells at low density into syngeneic BALB/c mice where CD69-high cells more efficiently formed tumors (**Fig. 6G-H**), with unenriched cells representing the median of these two conditions (**fig. S21K-P**). There was also a trend toward increased lung metastasis in mice implanted with CD69-high 4T1 cells (**fig. S21Q-R**). Magnetic sorting and CRISPR-a experiments therefore jointly established that neoplastic CD69 impacts early tumor growth.

**Figure 6.**
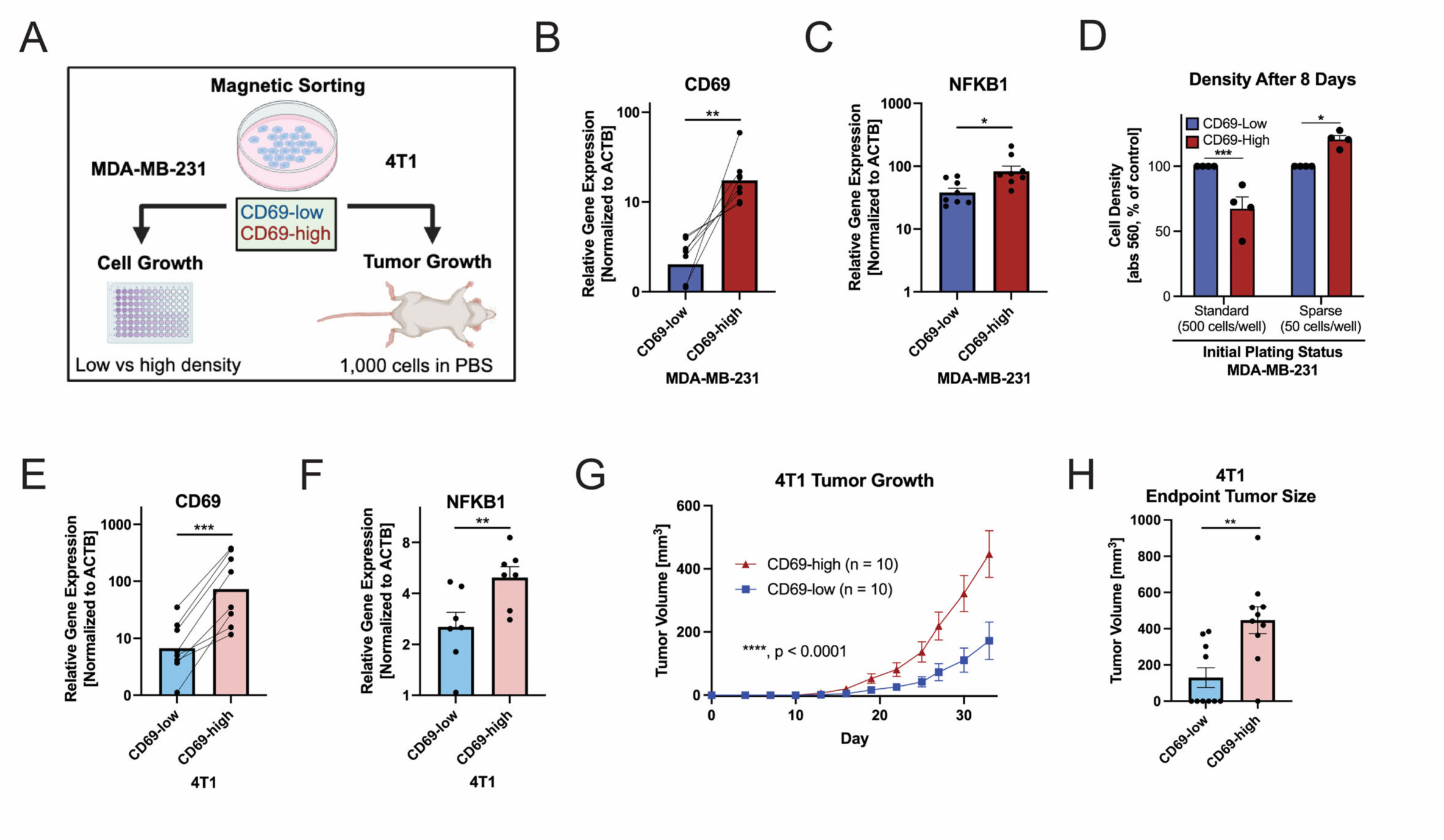
MDA-MB-231 and 4T1 cells magnetically sorted for CD69 grow more efficiently at low density *in vitro* and in mice. (**A**) MDA-MB-231 and 4T1 cells were magnetically sorted into CD69-low and CD69-high fractions. Sorting 200 million cells from each cell line yielded, on average, ∼50,000 CD69-high cells that were subjected to enrichment validation via qRT-PCR. MDA-MB-231 sorted populations were used for *in vitro* growth assays whereas 4T1 populations were assessed for tumor growth at low density in syngeneic BALB/c mice. MDA-MB-231 cells were enriched for (**B**) CD69 and (**C**) NFKB1 RNA expression. Data reflect eight biological replicates. (**D**) Subjecting these CD69-sorted MDA-MB-231 subpopulations to an *in vitro* growth assay revealed enhanced density by the CD69-high fraction when plated sparsely. Data reflect four biological replicates. (**E**) The 4T1 CD69-high fraction was enriched for CD69 and (**F**) NFKB1 RNA expression. Data reflect seven biological replicates. (**G**) CD69-high 4T1 cells more efficiently form tumors when injected into BALB/c mice at low density with (**H**) tumors reaching a larger size at endpoint. * p < 0.05; ** p < 0.01; *** p <0.001; **** p < 0.0001. 10 mice per group. Student’s t-test [(B), (C), (E), (F) and (H)], or two-way ANOVA (G) with Šidák correction (D). Data are means ± SEM.

We subsequently conducted loss-of-function studies to determine the extent to which neoplastic CD69 governed early tumors and hypothesized its absence would be detrimental to growth kinetics. These experiments were conducted in MMTV-PyMT-chOVA transgenic mice (43) to assess how loss of epithelial CD69 impacted the kinetics of tumor progression. Before initiating these studies, we confirmed that MMTV-PyMT-chOVA neoplastic mammary epithelial cells contained a CD69-positive subpopulation (**Fig. 7A**) expressing higher levels of Ki67 (**Fig. 7B**), and then subsequently intercrossed MMTV-PyMT-chOVA mice with mice harboring a homozygous deletion of CD69 (44,45) (**Fig. 7C; fig. S22A**). Tumor incidence in PyMT-CD69^KO/KO^ mice was not significantly altered from the PyMT-CD69^KO/+^ control (**fig. S22B-D**), however, early tumor growth was significantly delayed in PyMT-CD69^KO/KO^ mice where individual tumors exhibited an increased doubling time corresponding with a lower growth rate (**Fig. 7D-G**). As such, total tumor burden was decreased in PyMT-CD69^KO/KO^ mice (**Fig. 7H)** despite on average developing similar numbers of tumors as the PyMT-CD69^KO/+^ control (**fig. S22E**).

**Figure 7.**
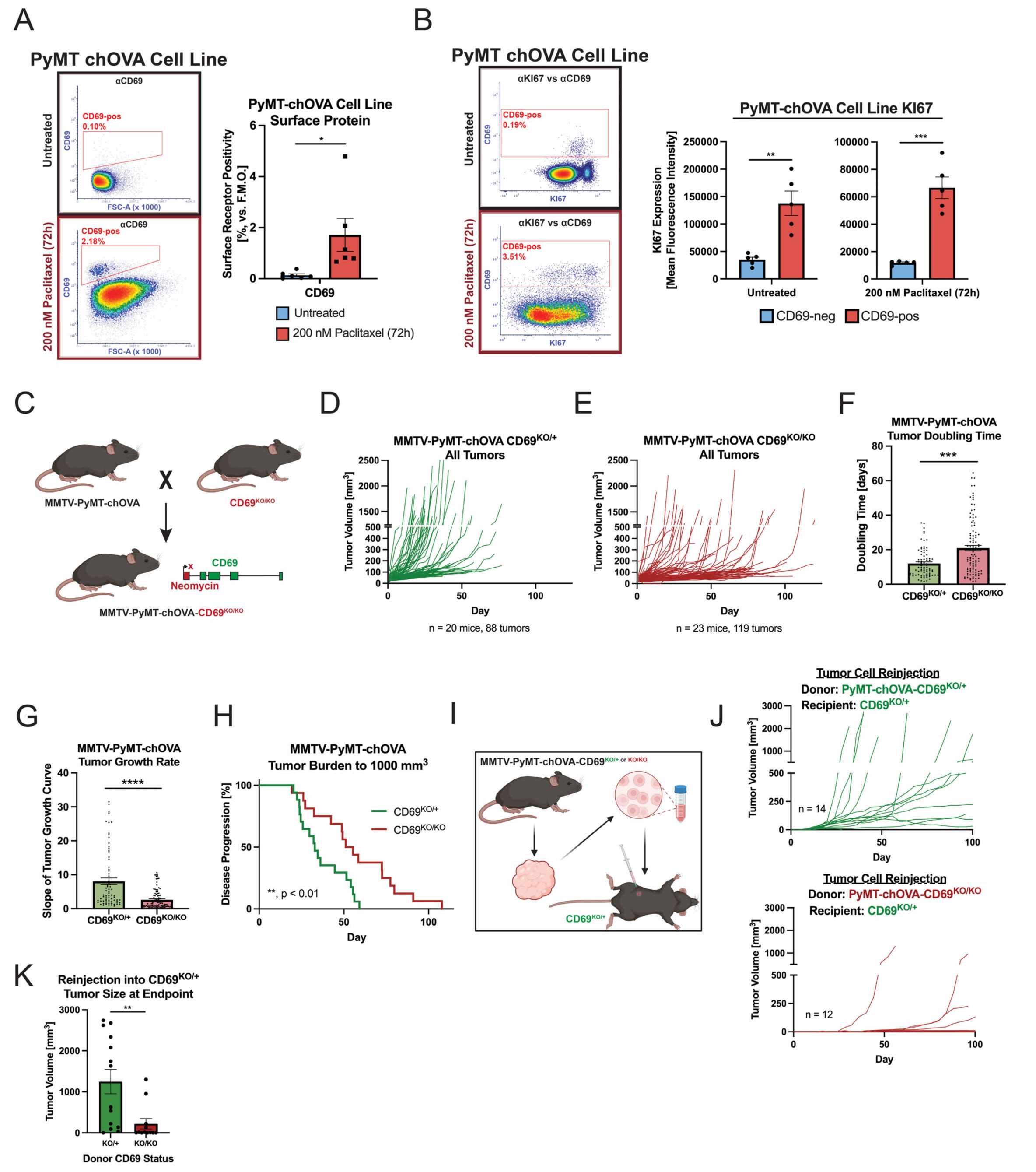
Knockout of CD69 in MMTV-PyMT-chOVA mice attenuates early tumor growth. (**A**) A cell line derived from an MMTV-PyMT-chOVA tumor contains a CD69-pos subpopulation that expands upon paclitaxel treatment and (**B**) upregulates Ki67. Data reflect six or five biological replicates, respectively. (**C**) MMTV-PyMT-chOVA mice were bred with CD69-homozygous null (ko/ko) mice to generate MMTV-PyMT-chOVA-CD69^KO/KO^ mice. (**D-E**) Comparing spontaneous tumor growth in PyMT-chOVA-CD69^KO/+^ versus MMTV-PyMT-chOVA-CD69^KO/KO^ mice uncovers (**F**) increased tumor doubling time and (**G**) slower rate in the latter. PyMT-chOVA-CD69^KO/+^, n = 88 tumors in 20 mice. PyMT-chOVA-CD69^KO/KO^, n = 119 tumors in 23 mice. Individual tumors depicted in (F) and (G). (**H**) MMTV-PyMT-chOVA-CD69^KO/KO^ mice take longer to achieve a tumor burden of 1000 mm^3^ when compared to mice harboring intact CD69. (**I**) Tumor cells were dissociated from MMTV-PyMT-chOVA-CD69^KO/+^ and MMTV-PyMT-chOVA-CD69^KO/KO^ mice for orthotopic reinjection back into standard CD69^KO/+^ mice. (**J**) MMTV-PyMT-chOVA-CD69^KO/+^ tumor cells are better able to grow upon reinjection into CD69^KO/+^ mice and (**K**) attain larger tumors by study endpoint. PyMT-chOVA-CD69^KO/+^ reinjection, n = 14 mice. PyMT-chOVA-CD69^KO/KO^ reinjection, n = 12 mice. * p < 0.05; ** p < 0.01; *** p <0.001; **** p < 0.0001. Student’s t-test [(A), (B) and (K)], Mann-Whitney test [(F) and (G)], or Mantel-Cox log-rank test (H). Data are means ± SEM.

We hypothesized that differences in host-derived stromal CD69 could underlie some of the *in vivo* tumor growth phenotype, thus we dissociated PyMT-CD69^KO/+^ and PyMT-CD69^KO/KO^ tumor cells and injected them back into syngeneic CD69-proficient (CD69^KO/+^) control mice (**Fig. 7I**). We selected three pairs of tumors for this experiment that arose in equivalent mammary glands and reached similar endpoint sizes (**fig. S22F-H**). The attenuated ability of dissociated PyMT-CD69^KO/KO^ cells to reestablish tumors in CD69^KO/+^ recipients demonstrated retention of their original phenotype (**Fig. 7J-K**) and recapitulated observations using magnetically-sorted 4T1 cells (**Fig. 6H**). Additional support for CD69 intrinsically regulating neoplastic growth was provided by CRISPR/Cas9 experiments with MDA-MB-231 cells (**fig. S22I**), where CD69 disruption prevented CD69 induction and attenuated cell growth *in vitro* (**fig. S22J-K**). We ultimately conclude that neoplastic breast cancer cells express leukocyte surface receptors and neoplastic CD69, in particular, potentiates malignant growth amid stress *in vitro* and in tumors and metastases *in vivo*.

## Discussion

Our study uncovered pervasive neoplastic immune mimicry in human breast tumors and related cell lines. In public human breast tumor scRNA-seq datasets, a minority subpopulation of neoplastic epithelial cells expressed combinations of 23 surface receptor genes typically attributed to leukocytes including CD3, CD14, CD18, CD45, CD68, CD69, CD74, and CD83. Neoplastic cells engaging in immune mimicry concomitantly upregulated dedifferentiation programs, NFκB pathway and proliferative signatures. Additionally, we detected rare subpopulations of presumed non-neoplastic epithelia expressing leukocyte surface receptors in reduction mammoplasties. Neoplastic CD69 was elevated in breast tumors versus reduction mammoplasties, especially those treated with neoadjuvant cytotoxic therapy and coexpressed in cells with a proliferative phenotype. We next validated epithelial CD69 in a histopathological cohort of 58 humor breast tumors by multiplexed IHC. Our research also revealed consistent, albeit small, subpopulations expressing canonical immune surface receptors in breast and mammary cancer cell lines. Neoplastic immune mimicry was further induced in aggressive/metastatic cell lines subjected to cytotoxic chemotherapy, exemplified by MDA-MB-231 and 4T1. CD69-positive MDA-MB-231 and 4T1 cells were additionally enriched for Ki67 and potentiated early tumor growth upon implantation into mice. In MMTV-PyMT-chOVA mice, homozygous deletion of CD69 conversely hindered mammary tumor growth but future studies might consider interchanging CD69^KO/+^ and CD69^KO/KO^ bone marrow to better clarify involvement of stromal CD69 in this phenotype. More studies on neoplastic immune mimicry are also needed to determine how it influences distal metastases, progression-free survival, overall survival, and therapeutic response/resistance in human disease.

In leukocyte biology, CD69 is a prominent indicator of early T cell activation with multiple functions including facilitating tissue retention and cell metabolism (25,26). Being an early-activation indicator, CD69 is challenging to study due to its dynamic regulation in cells; a fact woven throughout our efforts to discern its functional relevance in breast cancer before potentially serving as an early biomarker for patient stratification. This concept is highlighted by our observation that neoplastic CD69 may be associated with increased cell proliferation, especially amid harsh conditions such as cytotoxic chemotherapy or in distant metastases. Recently, others have reported a role for epithelial CD69 in ovarian cancer that emerges under hypoxia (49), indicating yet another route for CD69 induction. Confirming elevated neoplastic CD69 in treated breast tumors by multiplexed IHC was also challenging due to patients varying extensively in tumor grade, disease subtype, and other demographic features within the cohort. Consequently, forthcoming clinical investigations should evaluate neoplastic CD69 status via longitudinal sampling to ascertain its baseline and then acquired range throughout a therapeutic regimen. Harnessing antibodies to deliver innovative therapies to CD69-positive neoplastic cells could provide a novel strategy for overcoming drug resistance if the subpopulation eludes growth control.

Although we sought to document convincing evidence for neoplastic immune mimicry in breast tumors and cell lines, we recognize that additional research is required to unveil the true extent and functional significance of this cell state. Several surface markers upregulated within immune-mimicked scRNA-seq clusters have already been documented in breast epithelia, with CD14, CD44, CD74, and CXCR4 being notable examples (*15, 18, 19*), yet others remain underexplored. A next step could entail probing DNA within cells to determine whether expression of canonical leukocyte receptors is associated with specific breast cancer mutations. Lineage-tracing or barcoding experiments, especially those conducted *in vivo*, could also help demonstrate the neoplastic origin of immune-mimicked cells, the scope of plasticity in expression of immune markers, and which markers correlate with survival outcomes. Further investigating transcriptional regulators beyond the NFκB pathway that might elicit leukocyte gene expression in neoplastic cells would likewise be informative, as would studies determining if the function of these cell surface receptors in neoplastic cells reflect their known roles in leukocytes. Because leukocyte gene expression in structural cells outside the immune system may modulate immune-epithelial interactions (50), future research should hence address whether immune-mimicked cells evade anti-tumoral immunity akin to neoplastic cells expressing leukocyte surface receptors in other cancer lineages (16,17,21).

Neoplastic cells generally expressing leukocyte surface receptors would have ramifications for the detection and treatment of breast cancer. For example, we have previously reported that leukocytes readily infiltrate breast tumors following cytotoxic therapy (51,52), yet here we also establish that neoplastic epithelia express immune markers including CD45 and CD69 in similar conditions. It is therefore possible that immune mimicry leads to neoplastic cells avoiding clinical detection after cytotoxic therapy, due to our current reliance on surface features when discerning cell types. Since no tumor samples in our study were subjected to immunotherapeutics, the degree to which that modality fosters immune mimicry remains an open question. Regardless, it has been reported that circulating tumor cells bound to CD45-positive cells described as leukocytes more effectively grow as distant metastases (53,54). In light of our research, it is possible that some of these assumed “leukocytes” could be neoplastic cells endogenously expressing CD45 when transiting systemically, a fact that, if confirmed, could inspire new approaches to detect micrometastases. These findings thematically intersect with previous research identifying CD45-positive neoplastic-immune hybrids in the blood of patients afflicted with uveal melanoma (20) and the biomarkers affiliated with these cells (55). Since our flow cytometry analyses revealed that neoplastic breast cells endogenously positive for CD45 are significantly smaller in size than their CD45-negative counterparts, we caution against utilizing cell size as a distinguishing feature for leukocytes. Additional research in this area will be required before neoplastic immune mimicry can be clinically translated.

Computational biologists have questioned the utility of 2D embeddings to capture biological complexity during scRNA-seq analysis (56) and distinct scRNA-seq datasets can vary with respect to patient selection, time to treatment, sample processing, etc. We leveraged multiple public datasets in this study in pursuit of generalizability across cohorts and our approach demonstrates that it is sometimes worth thinking beyond cluster location when inferring cell types. Our method of initially gating breast epithelia via marker expression in lieu of cluster location exposed provocative mimicked neoplastic cell states with dedifferentiated features: developmental mimicry, endothelial mimicry, immune mimicry, mesenchymal mimicry, and neuronal mimicry. Although here we pursued neoplastic immune mimicry, the repertoire of genes that constitute each of these cell states and their roles in other cancer lineages deserves greater attention. Future studies might also consider if neoplastic cells engaging in these phenotypes could concomitantly lack mammary cytokeratins after, for example, an epithelial-mesenchymal transition (12). In the era of single-cell technologies, it is incumbent upon investigators to acknowledge and evaluate cell plasticity when conducting analyses.

## Acknowledgments

We dedicate this study to the memory of Dr. Zena Werb, who supported the early stages of this project prior to her unexpected death. In many ways, Zena was the idealized scientist – she was intelligent, creative, generous and wise, and she truly cared about those around her. Zena’s memory endures through her example and by the indelible mark she left on colleagues and mentees. The authors also thank members of the Werb, Heiser and Coussens laboratories for critical feedback and insight. This research acknowledges the Knight Cancer Institute P30 CA069533 grant and its supported shared resources (Flow Cytometry & Monoclonal Antibody Shared Resource, BioLibrary, Histopathology Shared Resource, Transgenic Mouse Model Core) and the Department of Comparative Medicine. The OHSU Knight Cancer Institute SMMART clinical trials program provided technical assistance and research contributions to the PANNTHR study supported by GSK (NCT04481113) and Eli Lilly and Company, who reviewed a preliminary version of the manuscript prior to submission. This study was further enabled by the following funding sources: NIH/NCI K00 CA212132 and the Collins Medical Trust to E.B.B.; NIH/NCI T32 CA254888 to E.B.B. and N.L.C.; Jayne Koskinas Ted Giovanis Foundation for Health and Policy, Kuni Foundation and NIH research grant U54-CA209988 to L.M.H.; the Susan G Komen Foundation and the National Foundation for Cancer Research to L.M.C. Dr. Steve Jameson generously provided the CD69^KO/KO^ mice and illustrations were created with BioRender.com. The Jayne Koskinas Ted Giovanis Foundation for Health and Policy is a private foundation committed to critical funding of cancer research. The opinions, findings, conclusions or recommendations expressed in this material are those of the author(s) and not necessarily those of the Jayne Koskinas Ted Giovanis Foundation for Health and Policy or its respective directors, officers, or staff. The authors are solely responsible for final content and interpretation.

## Notes

### Competing Interest Statement

L.M.C. has received reagent support from Cell Signaling Technologies, Syndax Pharmaceuticals, Inc., ZielBio, Inc., and Hibercell, Inc.; holds sponsored research agreements with Prospect Creek Foundation, grant support from Susan G. Komen Foundation, National Foundation for Cancer Research, and the National Cancer Institute; is on the Advisory Board for Carisma Therapeutics, Inc., CytomX Therapeutics, Inc., Kineta, Inc., Hibercell, Inc., Cell Signaling Technologies, Inc., Alkermes, Inc., NextCure, Guardian Bio, Dispatch Biotherapeutics, AstraZeneca Partner of Choice Network (OHSU Site Leader), Genenta Sciences, Pio Therapeutics Pty Ltd., and Lustgarten Foundation for Pancreatic Cancer Research Therapeutics Working Group. G.B.M. has received support from AstraZeneca, Zentalis, and Nanostring; is on the advisory board for Amphista, Astex, AstraZeneca, BlueDot, Ellipses Pharmaceuticals, ImmunoMET, Leapfrog Bio, Bruker/Nanostring, Neophore, Nerviano, Nuvectis, Pangea, PDX Pharmaceuticals, Qureator, Rybodyne, Signalchem Lifesciences, Tarveda, Turbine, and Zentalis Pharmaceuticals; has a financial stake in Bluedot, Catena Pharmaceuticals, ImmunoMet, Nuvectis, RyboDyne, SignalChem Lifesciences, Tarveda, and Turbine.

### Summary of Updates

This upload includes the revised manuscript resulting from peer review. Additional figures can be found in the supplement and textual edits were made to the manuscript.

