## Supplementary Figures and Tables for "Neoplastic immune mimicry potentiates breast tumor progression"

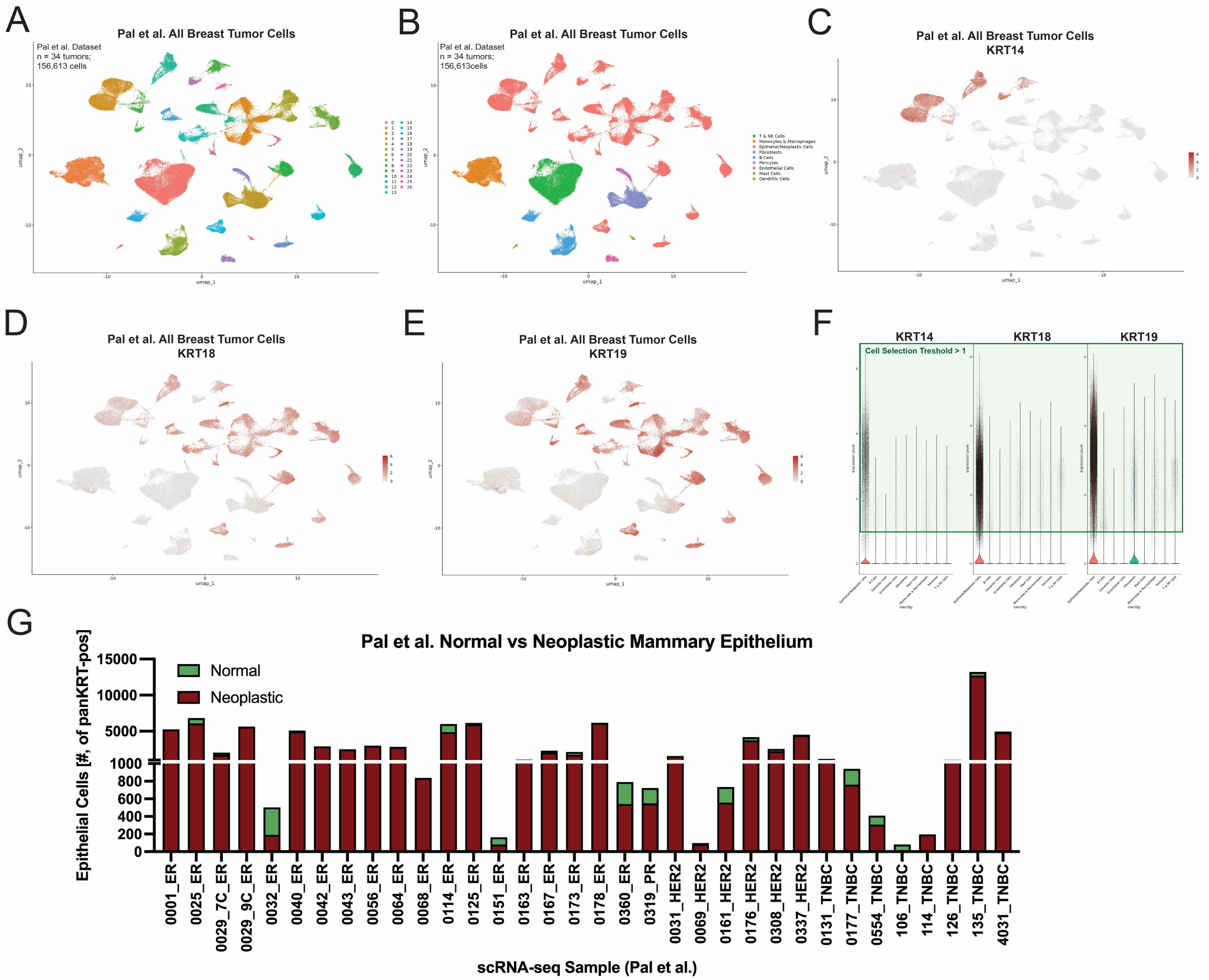


**Supplementary Figure 1**

**Supplementary Figure 1. Clusters of cells expressing mammary keratins and identification of the neoplastic compartment in the Pal *et al.* human breast tumor scRNA-seq dataset.** (**A**) Umap plot depicting all cells identified in the Pal *et al.* scRNA-seq dataset and (**B**) their conventional cell type assignments. Cells expressing mammary keratins (**C**) KRT14 (**D**) KRT18 and (**E**) KRT19 are located outside the main epithelial clusters. (**F**) Cells expressing mammary keratins were collected for downstream analysis regardless of cluster location using a threshold gating approach. (**G**) Neoplastic cells with inferred CNVs were refined from the epithelial pool. Normal epithelial cells showed parity with an epithelial reference generated from reduction mammoplasties and were hence omitted from downstream analysis.

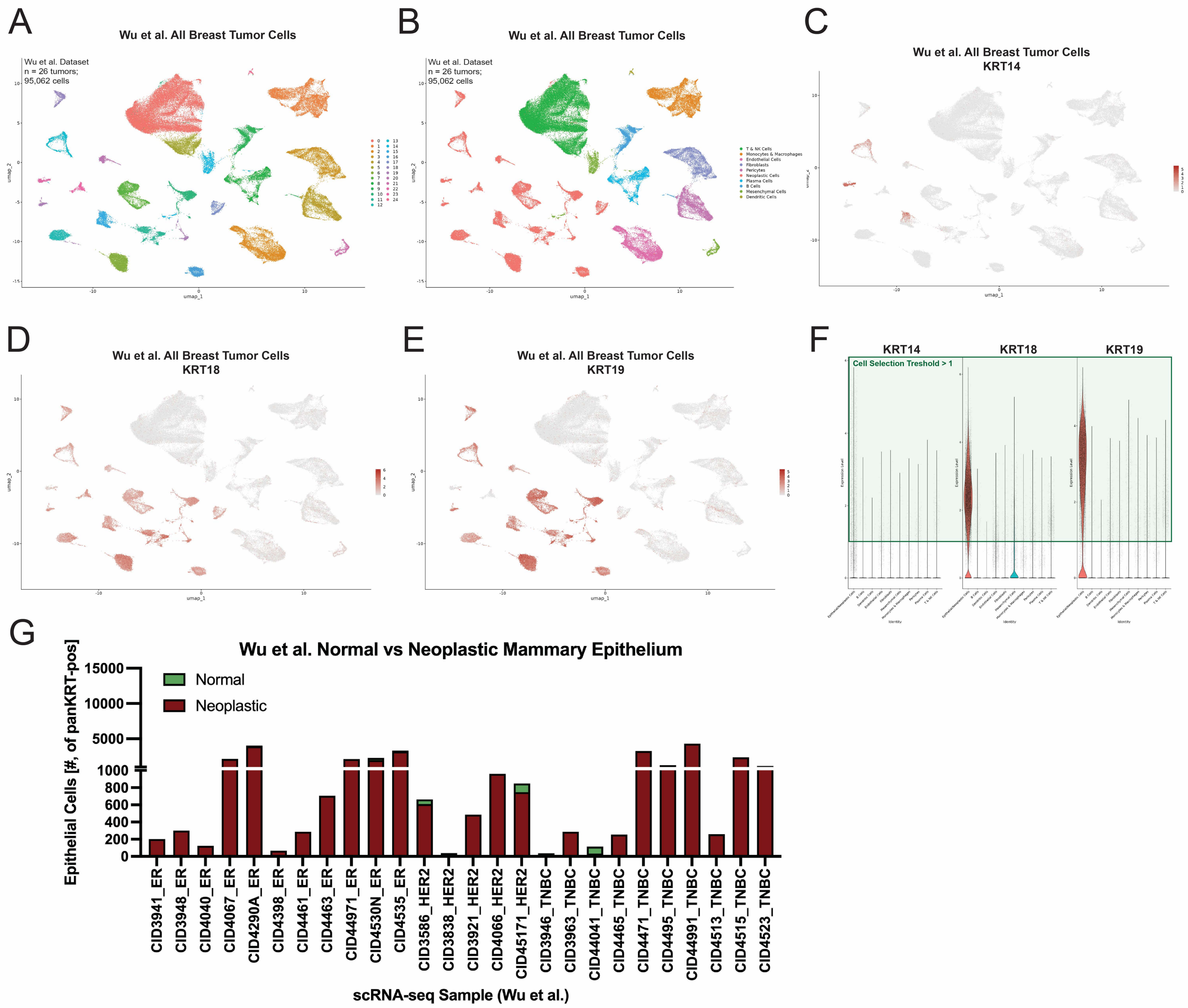


Supplementary Figure 2

**Supplementary Figure 2. Clusters of cells expressing mammary keratins and identification of the neoplastic compartment in the Wu *et al.* human breast tumor scRNA-seq dataset.** (**A**) Umap plot depicting all cells identified in the Wu *et al.* scRNA-seq dataset and (**B**) their conventional cell type assignments. Cells expressing mammary keratins (**C**) KRT14 (**D**) KRT18 and (**E**) KRT19 are located outside the main epithelial clusters. (**F**) Cells expressing mammary keratins were collected for downstream analysis regardless of cluster location using a threshold gating approach. (**G**) Neoplastic cells with inferred CNVs were refined from the epithelial pool. Normal epithelial cells showed parity with an epithelial reference generated from reduction mammoplasties and were hence omitted from downstream analysis.

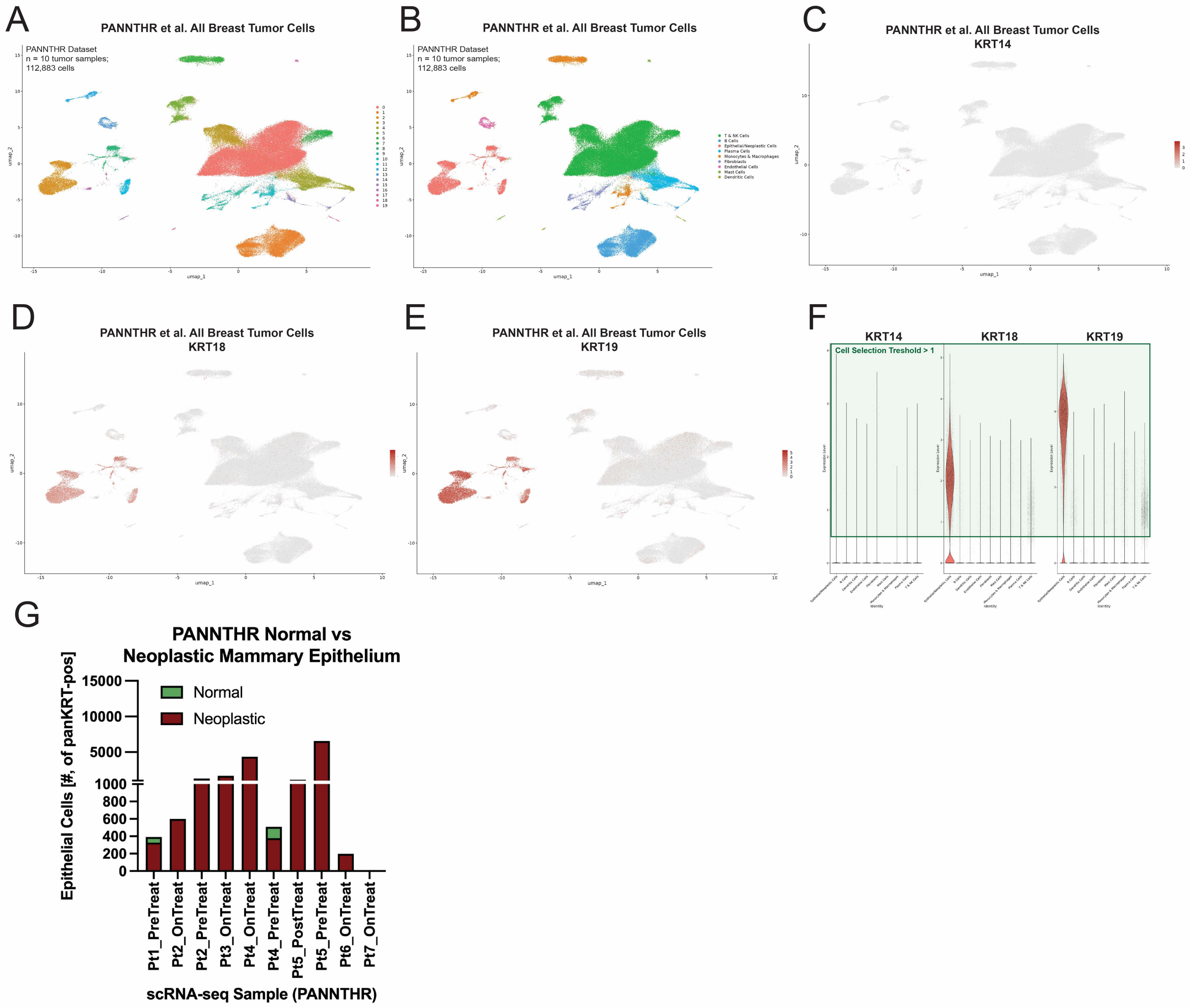


**Supplementary Figure 3**

**Supplementary Figure 3. Clusters of cells expressing mammary keratins and identification of the neoplastic compartment in the PANNTHR human breast tumor scRNA-seq dataset.** (**A**) Umap plot depicting all cells identified in the PANNTHR scRNA-seq dataset and (**B**) their conventional cell type assignments. Cells expressing mammary keratins (**C**) KRT14 (**D**) KRT18 and (**E**) KRT19 are located outside the main epithelial clusters. (**F**) Cells expressing mammary keratins were collected for downstream analysis regardless of cluster location using a threshold gating approach. (**G**) Neoplastic cells with inferred CNVs were refined from the epithelial pool. Normal epithelial cells showed parity with an epithelial reference generated from reduction mammoplasties and were hence omitted from downstream analysis.

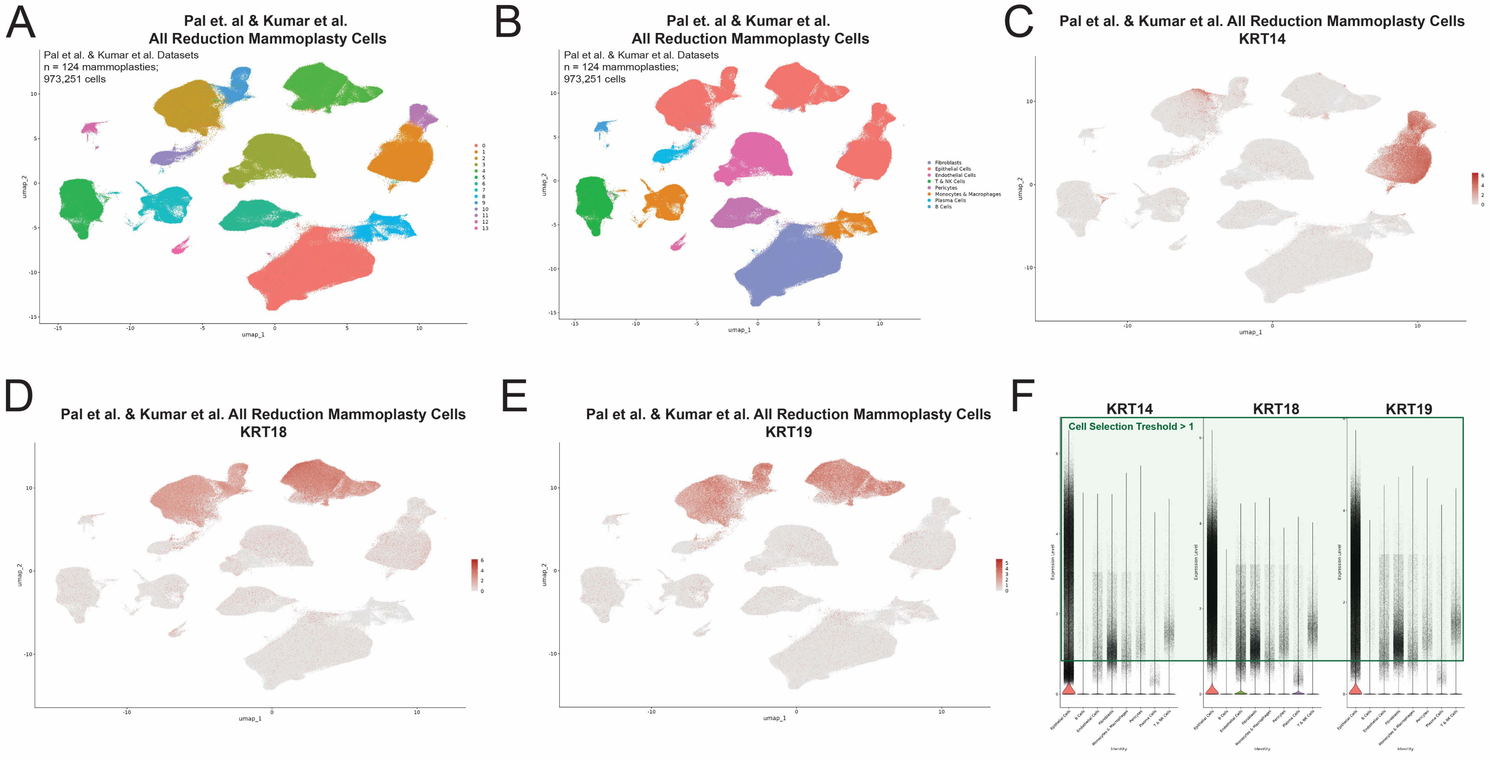


Supplementary Figure 4

**Supplementary Figure 4. Clusters of cells expressing mammary keratins and identification of mammary epithelial cells in reduction mammoplasty samples from the Pal *et al.* and Kumar *et al.* scRNA-seq datasets.** (**A**) Umap plot depicting all cells identified in the Pal *et al.* and Kumar *et al.* datasets and (**B**) their conventional cell type assignments. Cells expressing mammary keratins (**C**) KRT14 (**D**) KRT18 and (**E**) KRT19 are located outside the main epithelial clusters. (**F**) Cells expressing mammary keratins were collected for downstream analysis regardless of cluster location using a threshold gating approach.

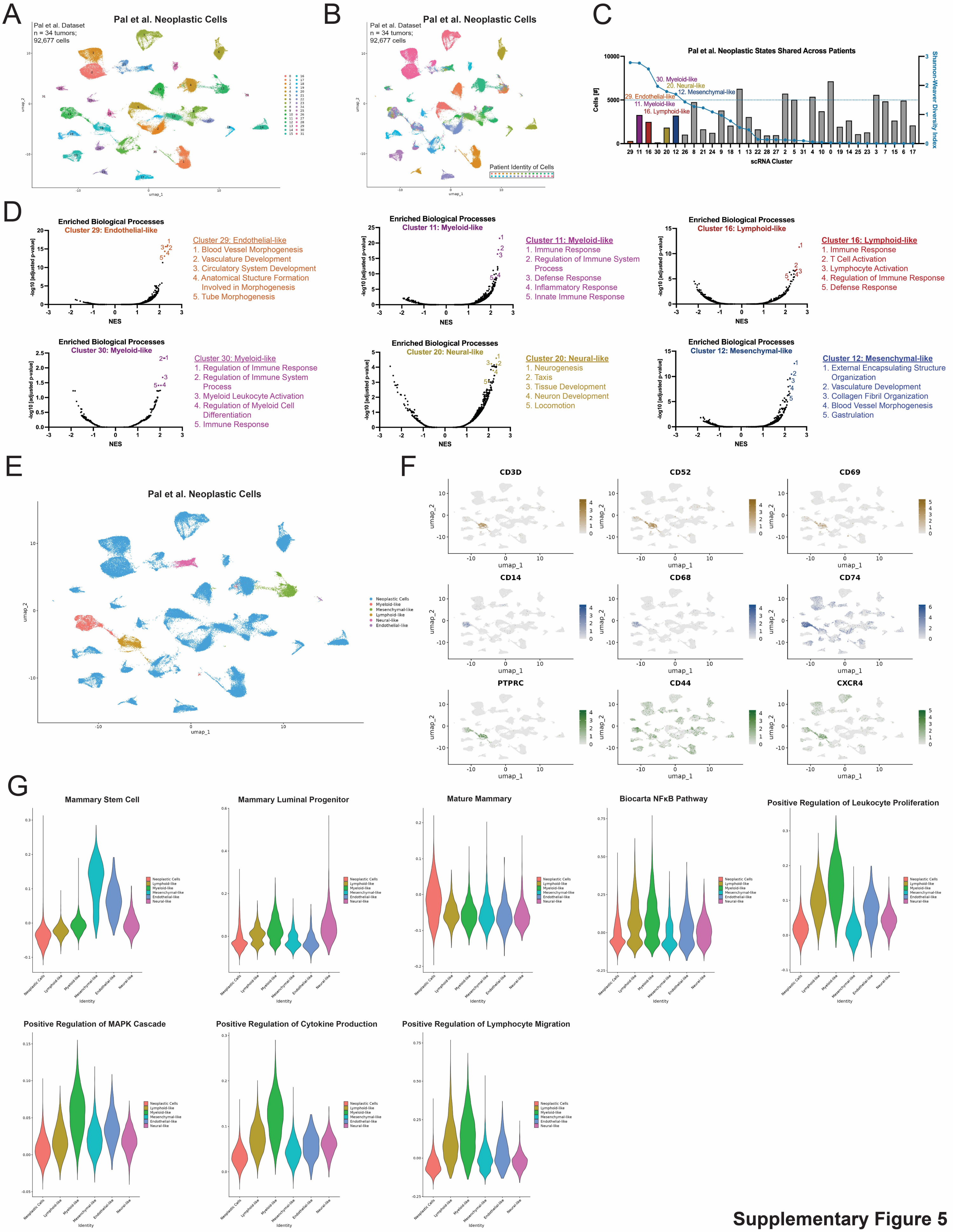


**Supplementary Figure 5**

**Supplementary Figure 5. Extended scRNA-seq analysis of mimicked cell states in the Pal *et al.* human breast tumor scRNA-seq dataset.** (**A**) Umap plot depicting neoplastic cells identified in the Pal *et al.* dataset and (**B**) the patient identity of those cells. (**C**) Neoplastic cells in each scRNA-seq cluster, their Shannon-Weaver Diversity Index, and their mimicry designation. (**D**) Mimicry designations are based on enriched Gene Ontology biological processes. (**E**) Umap plot with cluster annotations for mimicked cell states. (**F**) Expression of marker genes polarized toward lymphoid-like cells (gold), myeloid-like cells (blue) as well as both facets of immune mimicry (green). (**G**) The levels of key gene expression signatures for all mimicked cell states across the *Pal et al.* breast tumor dataset.

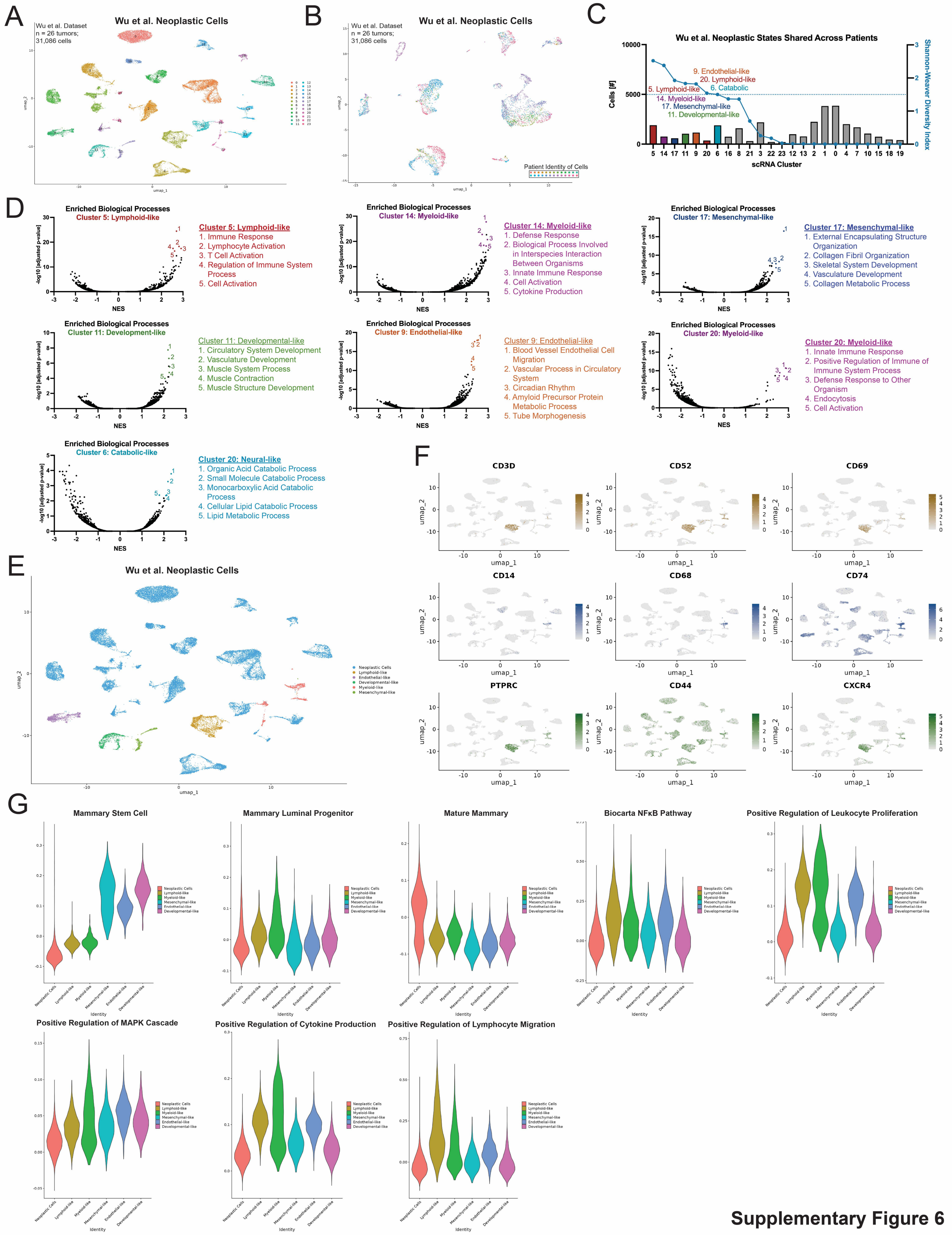


Supplementary Figure 6

**Supplementary Figure 6. Extended scRNA-seq analysis of mimicked cell states in the Wu *et al.* human breast tumor scRNA-seq dataset.** (**A**) Umap plot depicting neoplastic cells identified in the Wu *et al.* dataset and (**B**) the patient identity of those cells. (**C**) Neoplastic cells in each scRNA-seq cluster, their Shannon-Weaver Diversity Index, and their mimicry designation. (**D**) Mimicry designations are based on enriched Gene Ontology biological processes. (**E**) Umap plot with cluster annotations for mimicked cell states. (**F**) Expression of marker genes polarized toward lymphoid-like cells (gold), myeloid-like cells (blue) as well as both facets of immune mimicry (green). (**G**) The levels of key gene expression signatures for all mimicked cell states across the *Wu et al.* breast tumor dataset.

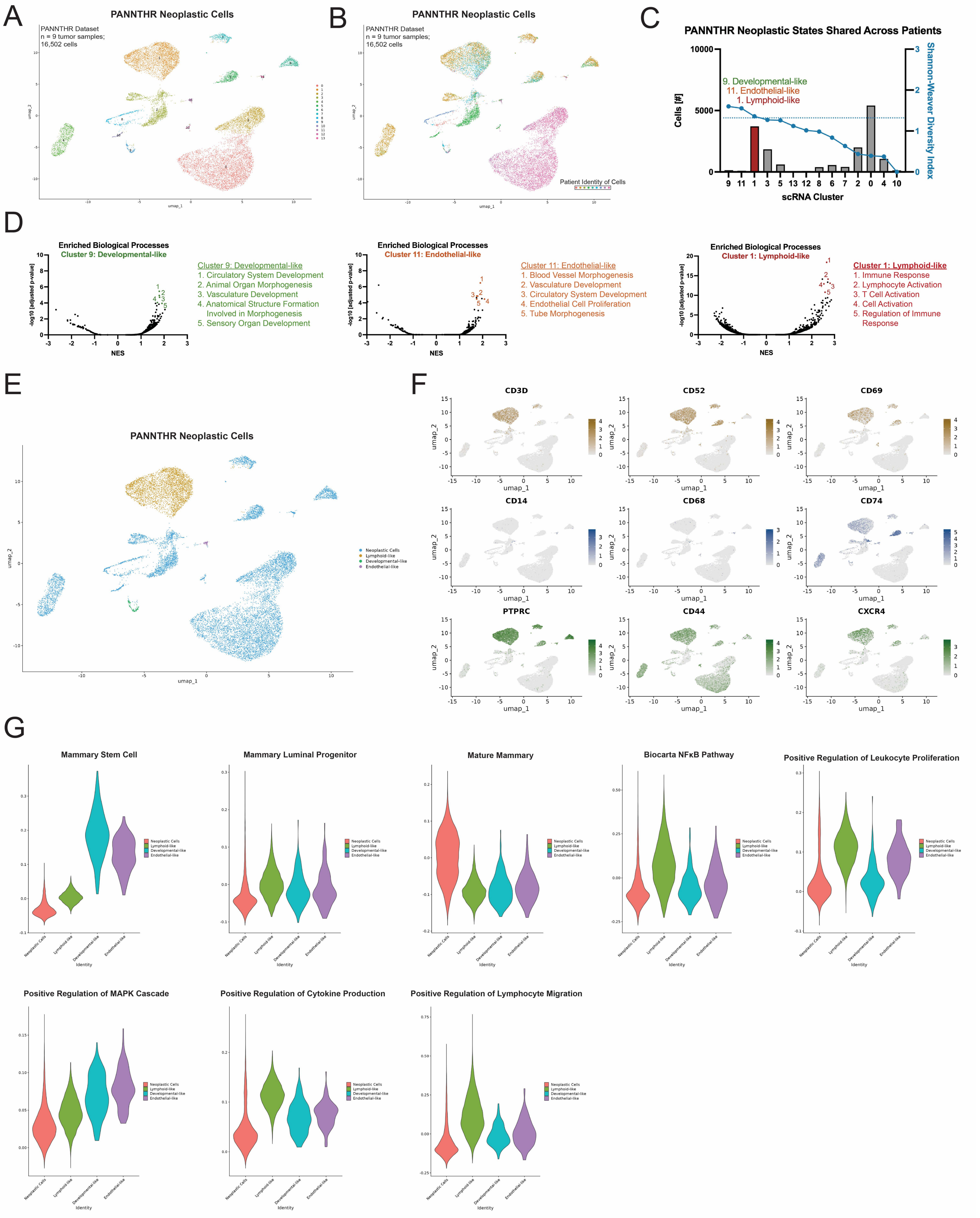


Supplementary Figure 7

**Supplementary Figure 7. Extended scRNA-seq analysis of mimicked cell states in the PANNTHR human breast tumor scRNA-seq dataset.** (**A**) Umap plot depicting neoplastic cells identified in the PANNTHR dataset and (**B**) the patient identity of those cells. (**C**) Neoplastic cells in each scRNA-seq cluster, their Shannon-Weaver Diversity Index, and their mimicry designation. Shannon-Weaver Diversity Index threshold lowered slightly to allow for collection of immune-mimicked cells. (**D**) Mimicry designations are based on enriched Gene Ontology biological processes. (**E**) Umap plot with cluster annotations for mimicked cell states. (**F**) Expression of marker genes polarized toward lymphoid-like cells (gold), myeloid-like cells (blue) as well as both facets of immune mimicry (green). (**G**) The levels of key gene expression signatures for all mimicked cell states across the PANNTHR breast tumor dataset.

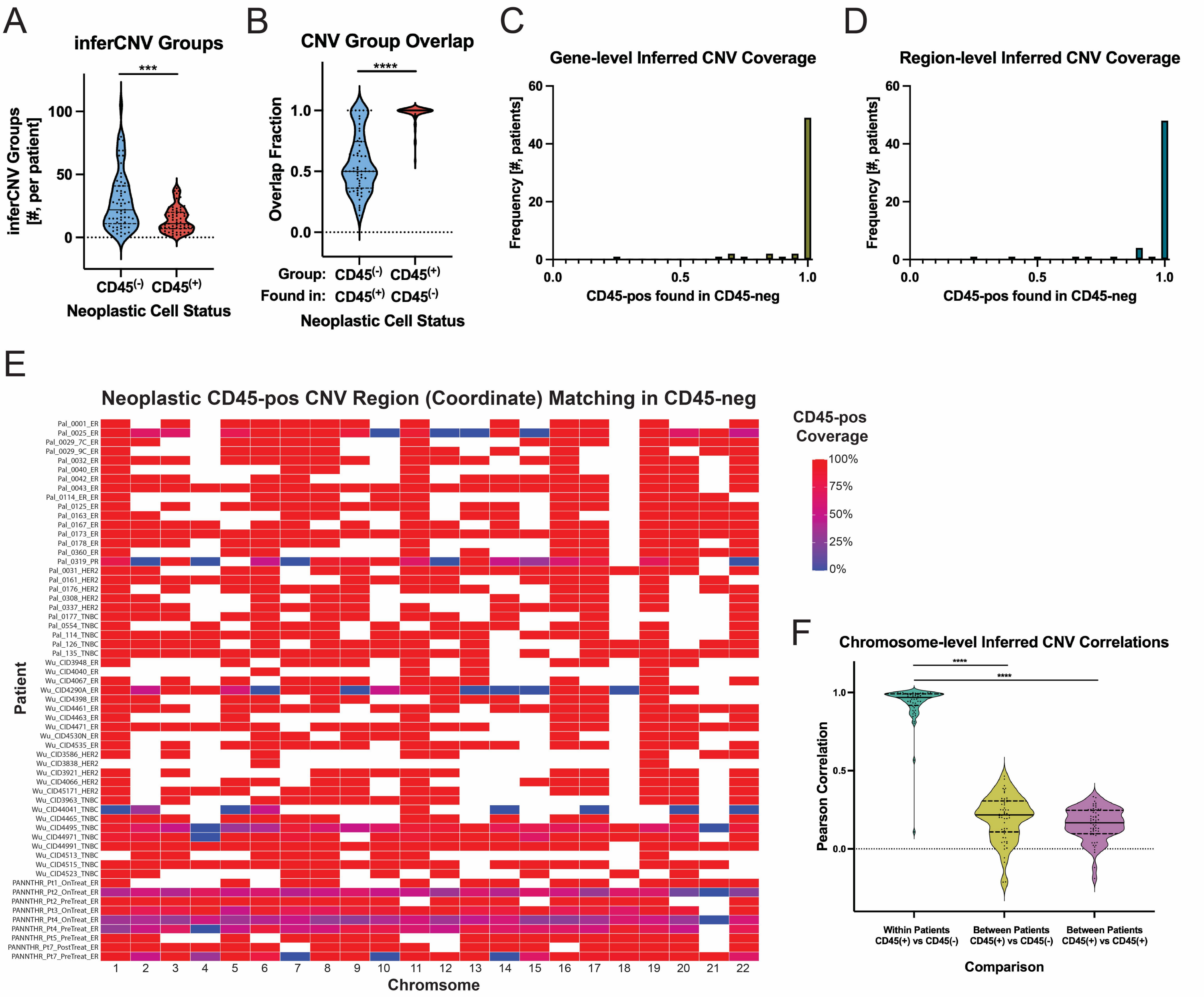


Supplementary Figure 8

**Supplemental Figure 8. CD45-pos (PTPRC-pos) cells with epithelial features show inferred CNVs from CD45-neg (PTPRC-neg) neoplastic cells by scRNA-seq.** (**A**) CD45-pos epithelial cells were represented by fewer inferCNV groups that (**B**) overlapped with the CD45-neg neoplastic population, implying CD45-pos enrichment from CD45-neg neoplastic epithelium. (**C**) Individual genes attributed to amplifications or deletions within inferred CNVs in CD45-pos cells are covered by CD45-neg cells on a patient-by-patient basis. (**D-E**) The exact genomic coordinates of discrete inferred CNVs from each patient's entire CD45-pos epithelial population were highly consistent with the same patient's CD45-neg population either (**D**) aggregated or (**E**) at the chromosome-level. (**F**) Chromosomes harboring inferred CNVs from CD45-pos epithelial cells were highly correlated with the CD45-neg neoplastic population within patients versus CD45-neg or CD45-pos populations between patients. *** $p < 0.001$; **** $p < 0.0001$. Mann-Whitney test [(A) and (B)] with means and quartiles depicted. Analyses performed on tumors containing at least 10 CD45-pos epithelial cells.

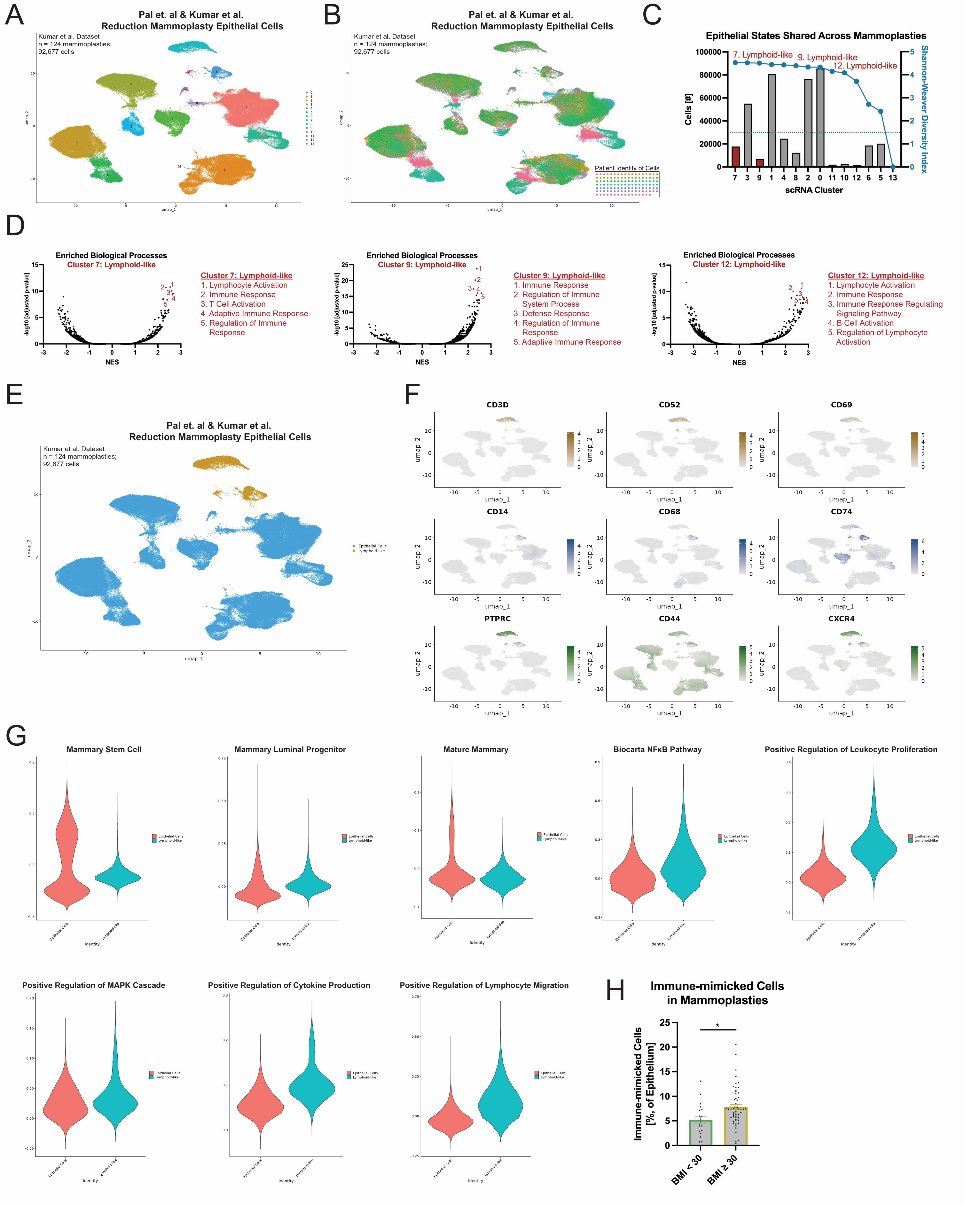


Supplementary Figure 9

**Supplementary Figure 9. Extended scRNA-seq analysis of mimicked cell states in reduction mammoplasties from the Pal *et al.* and Kumar *et al.* scRNA-seq datasets.** (**A**) Umap plot depicting neoplastic cells identified in the Pal *et al.* and Kumar *et al.* datasets and (**B**) the patient identity of those cells. (**C**) Neoplastic cells in each scRNA-seq cluster, their Shannon-Weaver Diversity Index, and their mimicry designation. (**D**) Mimicry designations are based on enriched Gene Ontology biological processes. (**E**) Umap plot with cluster annotations for mimicked cell states. (**F**) Expression of marker genes polarized toward lymphoid-like cells (gold), myeloid-like cells (blue) as well as both facets of immune mimicry (green). (**G**) The levels of key gene expression signatures for all mimicked cell states across the Pal *et al.* and Kumar *et al.* reduction mammoplasty datasets. (**H**) Immune-mimicked cells are more frequent in reduction mammoplasties from patients with a Body Mass Index (BMI) $\geq$ 30. Mammoplasties with BMI meta data, BMI $< 30$ (n = 20); BMI $\geq 30$ (n = 55). * $p < 0.05$; Student's t-test (H). Data are means ± SEM.

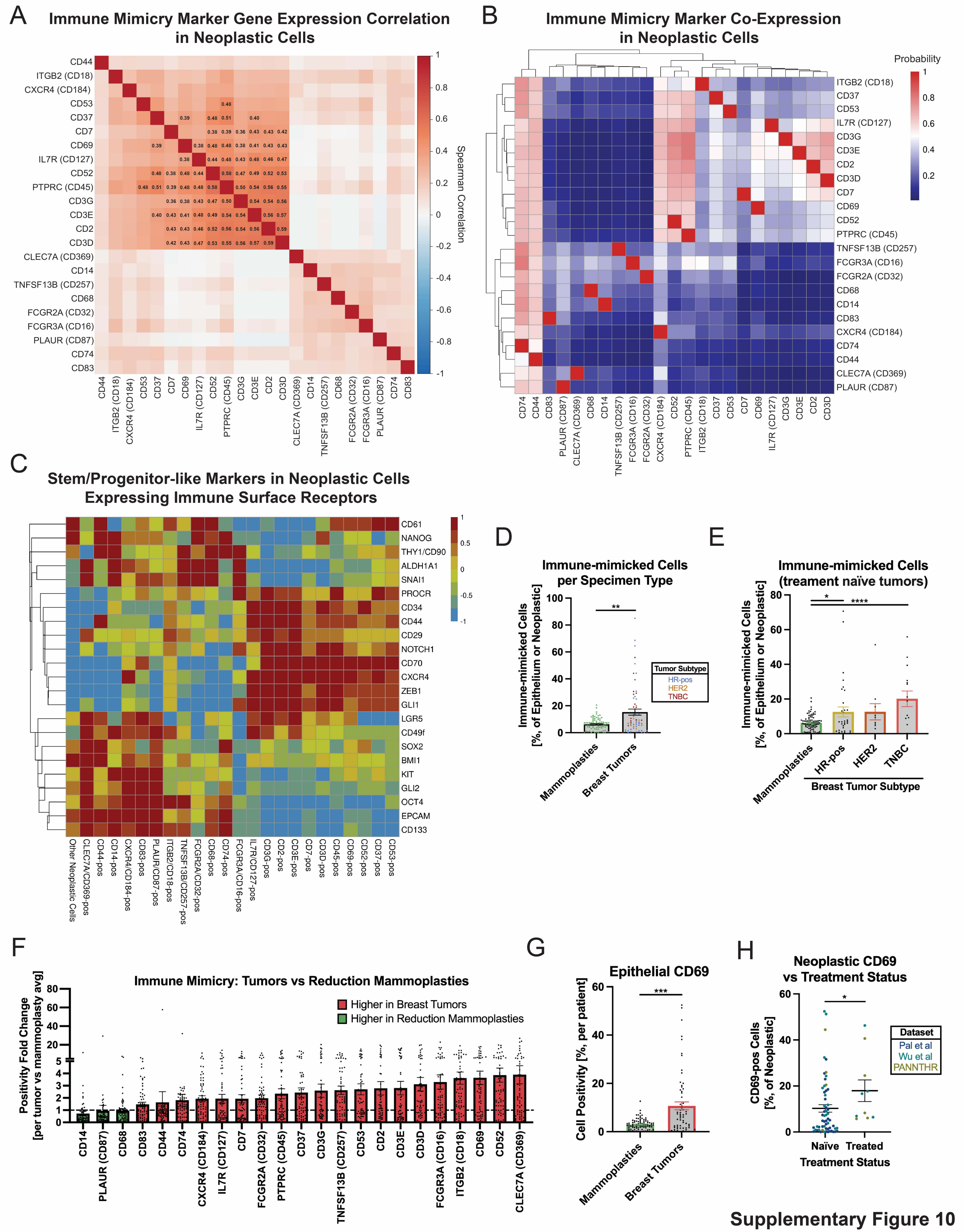


**Supplementary Figure 10**

**Supplementary Figure 10. Additional features of neoplastic immune mimicry identified in scRNA-seq datasets of human breast tumors and reduction mammoplasties.** (**A**) Gene expression correlation of immune mimicry surface markers across human breast tumors. Coefficients for significant correlations are indicated. (**B**) Co-expression of immune surface markers across human breast tumors. (**C**) Neoplastic expression of leukocyte surface receptor genes accompanies known tumor-initiation markers in breast cancer, with myeloid-like and lymphoid-like receptors being polarized toward different dedifferentiation signatures. (**D**) Neoplastic immune mimicry is elevated in breast tumors versus reduction mammoplasties (**E**) especially those from the triple-negative subtype. (**F**) Comparing the number of cells expressing each immune mimicry receptor in breast tumors versus reduction mammoplasties unveils CD69 to be one of the top markers associated with malignancy. (**G**) The number of neoplastic cells expressing CD69 is significantly elevated in breast tumors compared to reduction mammoplasties. (**H**) Neoplastic cells expressing CD69 are increased in tumors subjected to therapy before surgical resection. On this plot, patients from different scRNA-seq cohorts are indicated. * $p < 0.05$; ** $p < 0.01$; *** $p <0.001$; **** $p < 0.0001$. Spearman correlation (A), Mann-Whitney test [(D), (G) and (H)], or Kruskal-Wallis test (E). Data are means ± SEM.

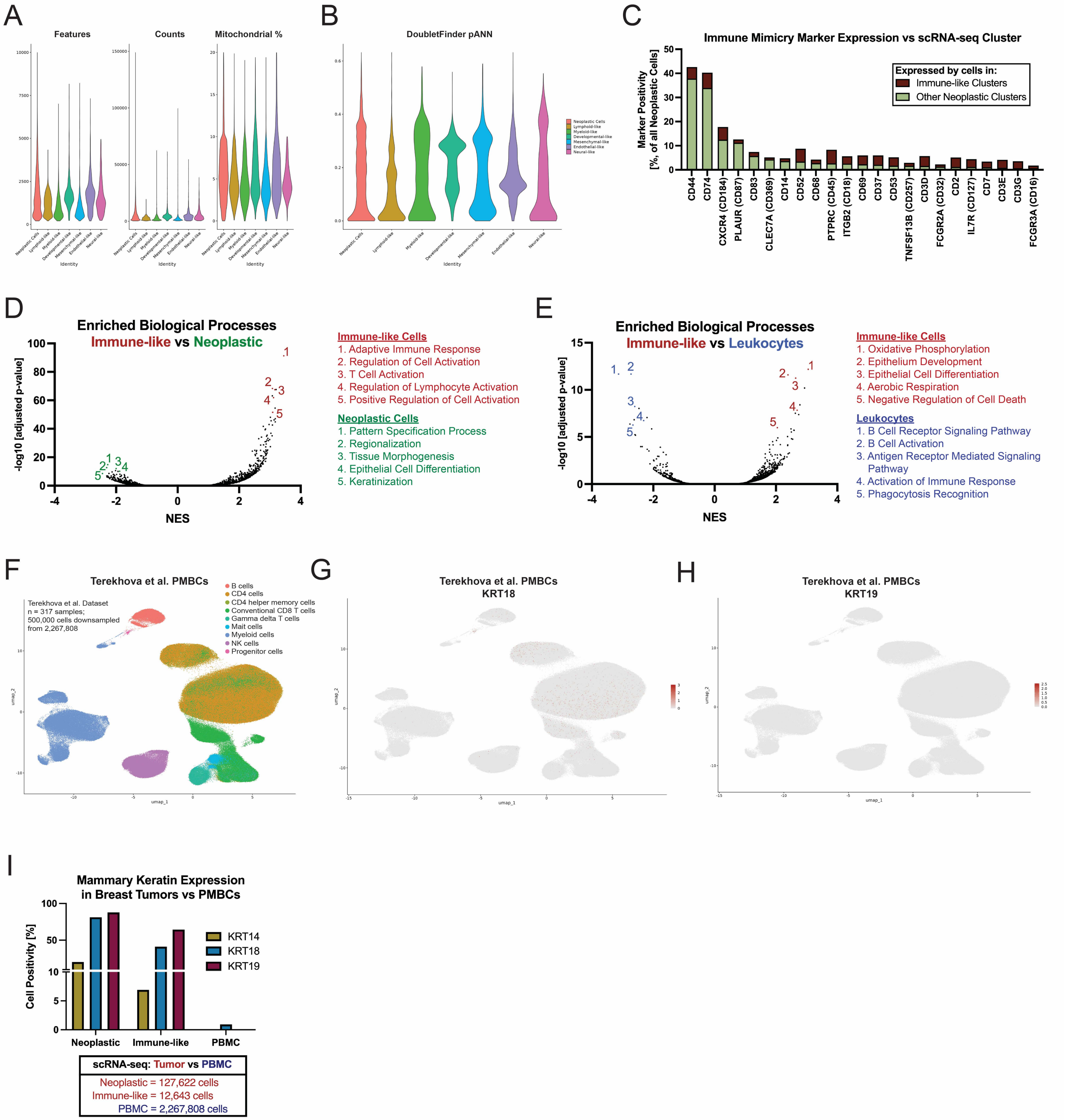


**Supplementary Figure 11**

**Supplementary Figure 11. Computational efforts to ascertain the origin of immune-mimicked cells.** (**A**) Assessment of the number of genes (features), molecules (RNA), and percent of mitochondrial reads in various neoplastic cell states found across human breast tumors. For each of these criteria, immune-mimicked cell states mirror that of non-mimicked neoplastic cells. (**B**) The proportion of DoubletFinder's artificial nearest neighbors for immune-mimicked cell states mirrors that of non-mimicked neoplastic cells. (**C**) Cells expressing many leukocyte surface receptors are found in both well-defined epithelial scRNA-seq clusters and immune-like clusters. (**D**) Immune-mimicked cells upregulated leukocyte biological processes when compared with non-mimicked neoplastic cells and (**E**) epithelial/growth processes when compared with tumoral leukocytes. (**F**) Analysis of over 2 million human peripheral blood mononuclear cells reveals (**G**) a tiny fraction expressing KRT18 yet virtually no expression of KRT14 or (**H**) KRT19. Umap shows 500,000 PMBCs downsampled from over 2 million. (**I**) The number of immune-like cells expressing KRT14, KRT18, and KRT19 generally mirrors that of non-mimicked neoplastic cells.

### Potential Neoplastic CD45 and/or CD68 in Human Breast Tumors

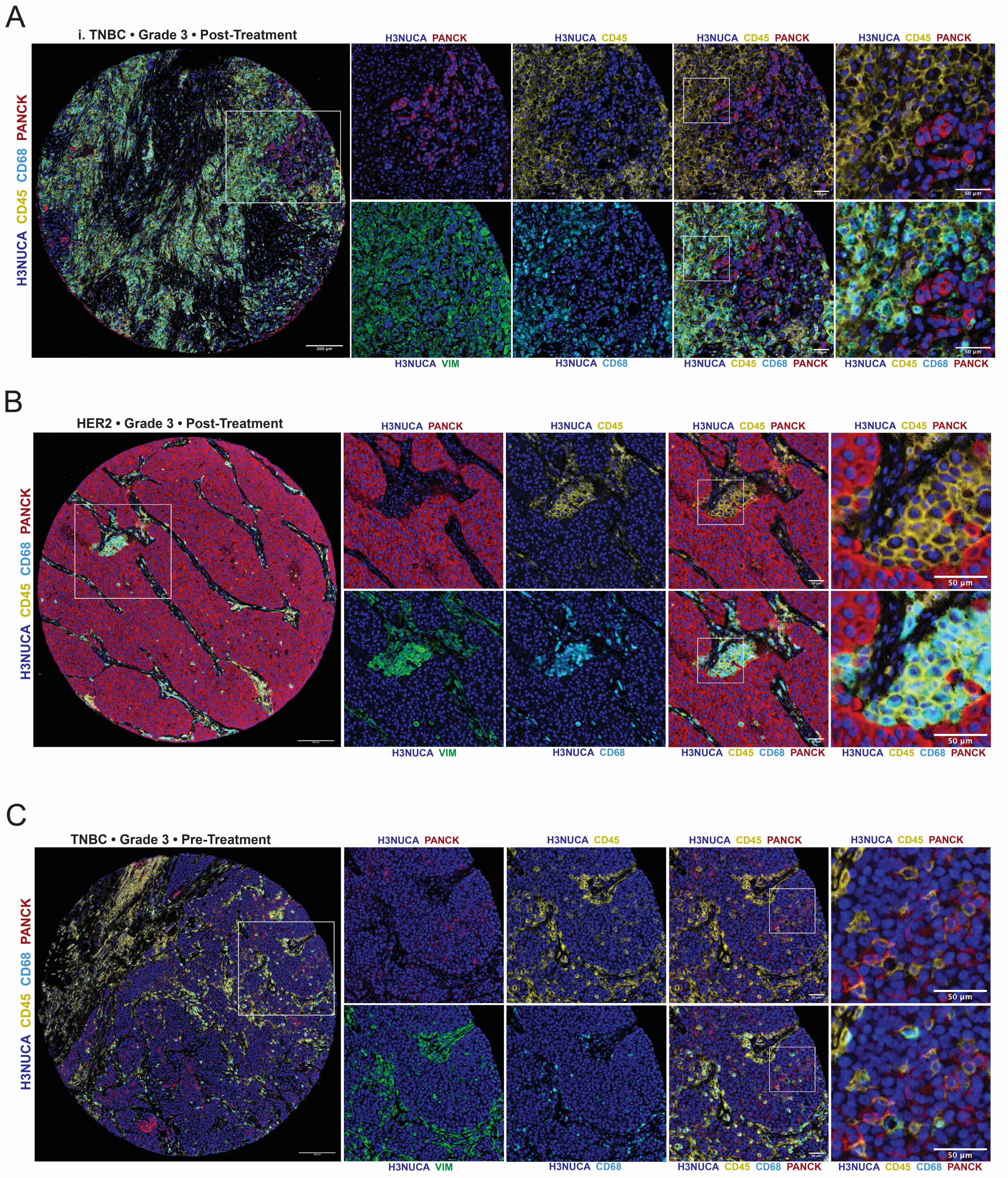


**Supplemental Figure 12**

**Supplemental Figure 12. Multiplexed immunohistochemistry examples of prospective CD45 and/or CD68 staining in neoplastic human breast epithelium.** (**A**) Possible neoplastic CD45 expression in a high-grade human triple-negative breast tumor, denoted by overlapping faint PANCK (red) and CD45 (yellow) cells. In this patient, a core of PANCK-pos cells also expressing vimentin (green) is surrounded by vimentin-pos and CD68-pos (cyan) cells that exhibit heterotypic junctions. This tumor region may indicate a zone of epithelial-mesenchymal transition. Scale bars, tissue core, 200 μm; inset, 50 μm. (**B**) Possible neoplastic CD45 expression in a high-grade HER2-positive breast tumor, denoted by overlapping PANCK (red) and CD45 (yellow) in cells near the tumor-stromal interface. These CD45-pos cells express vimentin (green) and CD68 (cyan) and display a morphological architecture that resembles adjacent PANCK-pos epithelium, suggestive of an epithelial-mesenchymal transition. A different region from this patient is depicted in Figure 2B (example ii.) showing epithelial CD69. Scale bars, tissue core, 200 μm; inset, 50 μm. (**C**) Possible neoplastic CD45 expression in a high-grade human triple-negative breast tumor, denoted by overlapping PANCK (red) and CD45 (yellow) in cells within tumor nests. Neoplastic epithelium in this patient does not strongly express PANCK or vimentin (green). Scale bars, tissue core, 200 μm; inset, 50 μm.

A

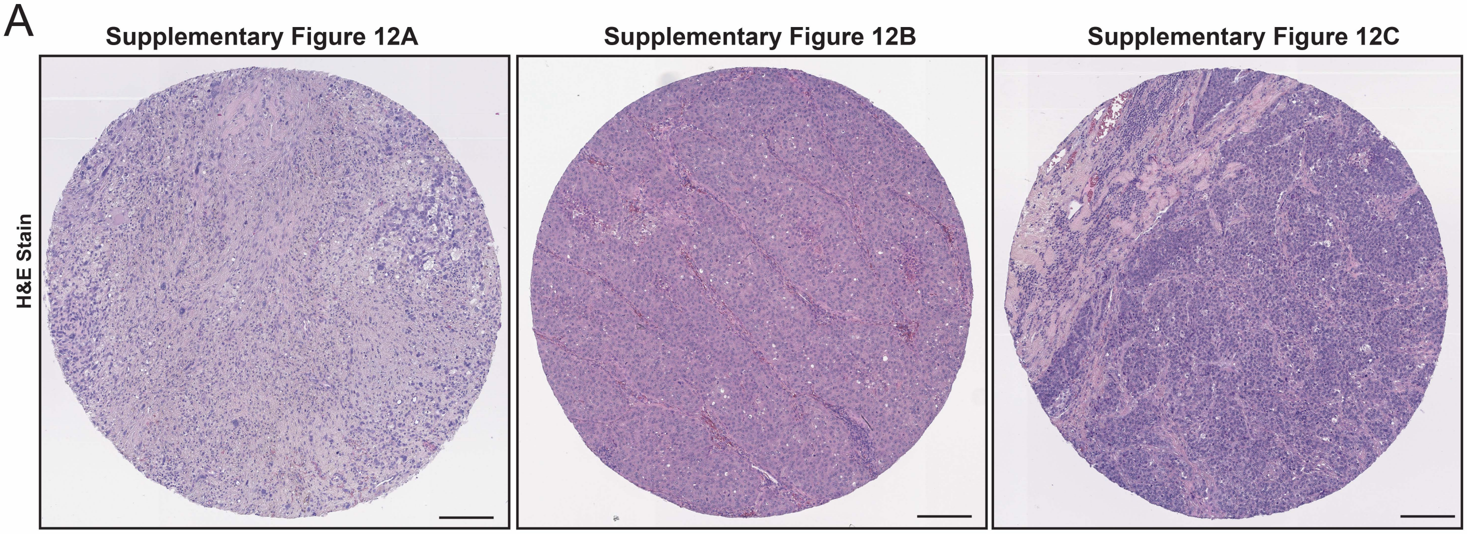


Supplemental Figure 13

**Supplemental Figure 13. Hematoxylin and eosin stains for example human breast tumor tissues.** (**A**) H&E stains of breast tumor cores depicted in Supplementary Figure 12. Scale bar, 200 μm. H&E staining was performed on tissue cores as the first step of the multiplexed immunohistochemistry workflow and thus preceded any antibodies.

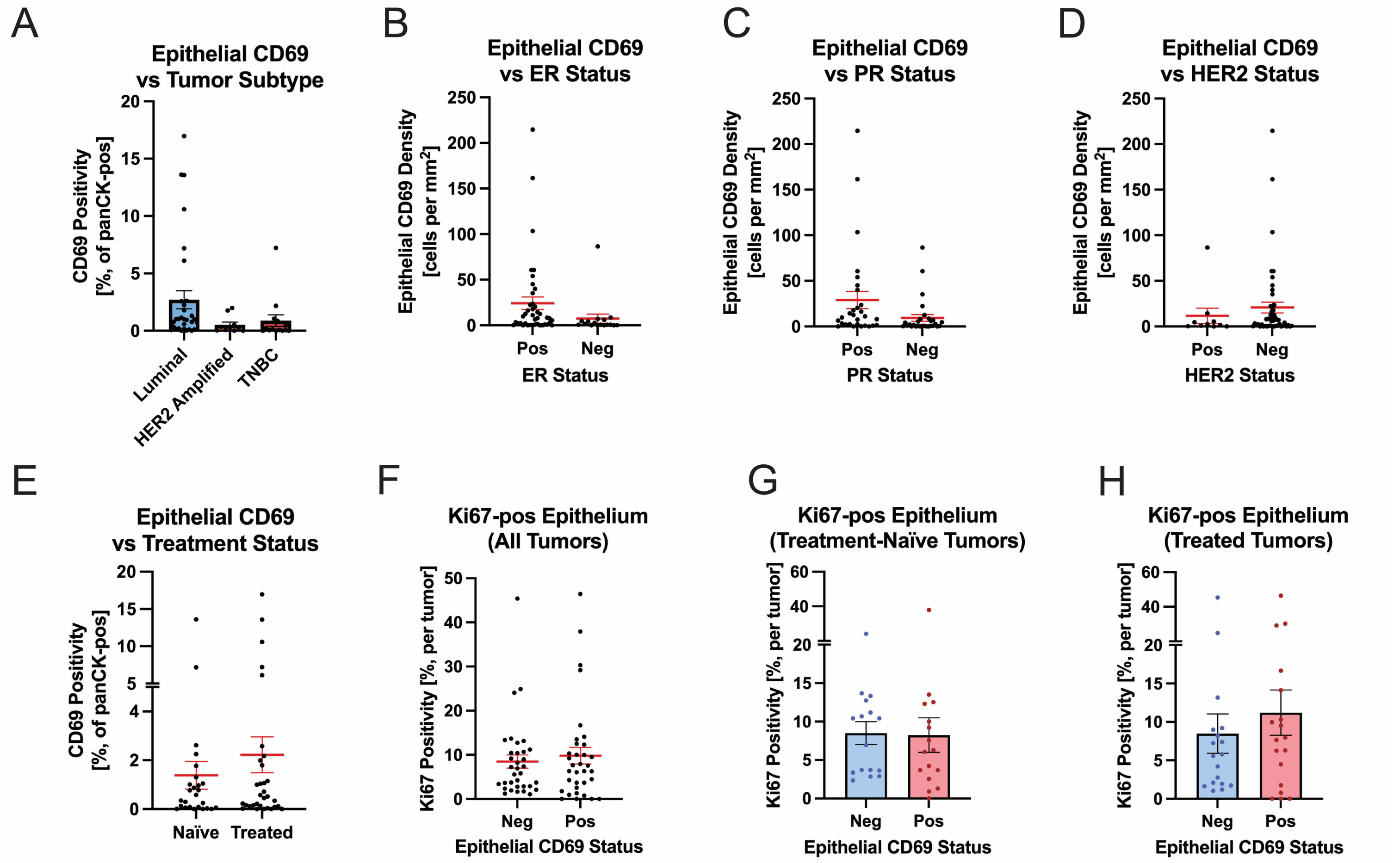


Supplementary Figure 14

**Supplementary Figure 14. Histopathologic epithelial CD69 in human breast tumors stratified by clinical parameters.** (**A**) The detection of epithelial CD69 in breast tumors according to their subtype. (**B-D**) Epithelial CD69 in individual tumors stratified by breast cancer receptor status such as the (**B**) estrogen receptor, (**C**) progesterone receptor, and (**D**) the HER2 receptor. (**E**) Tumors subjected to therapy before resection appear to have more epithelia expressing CD69. (**F-H**) Ki67 in CD69-neg versus CD69-pos epithelial cells across (**F**) all breast tumors, (**G**) treatment-naïve breast tumors, and (**H**) treated breast tumors. Data are means ± SEM.

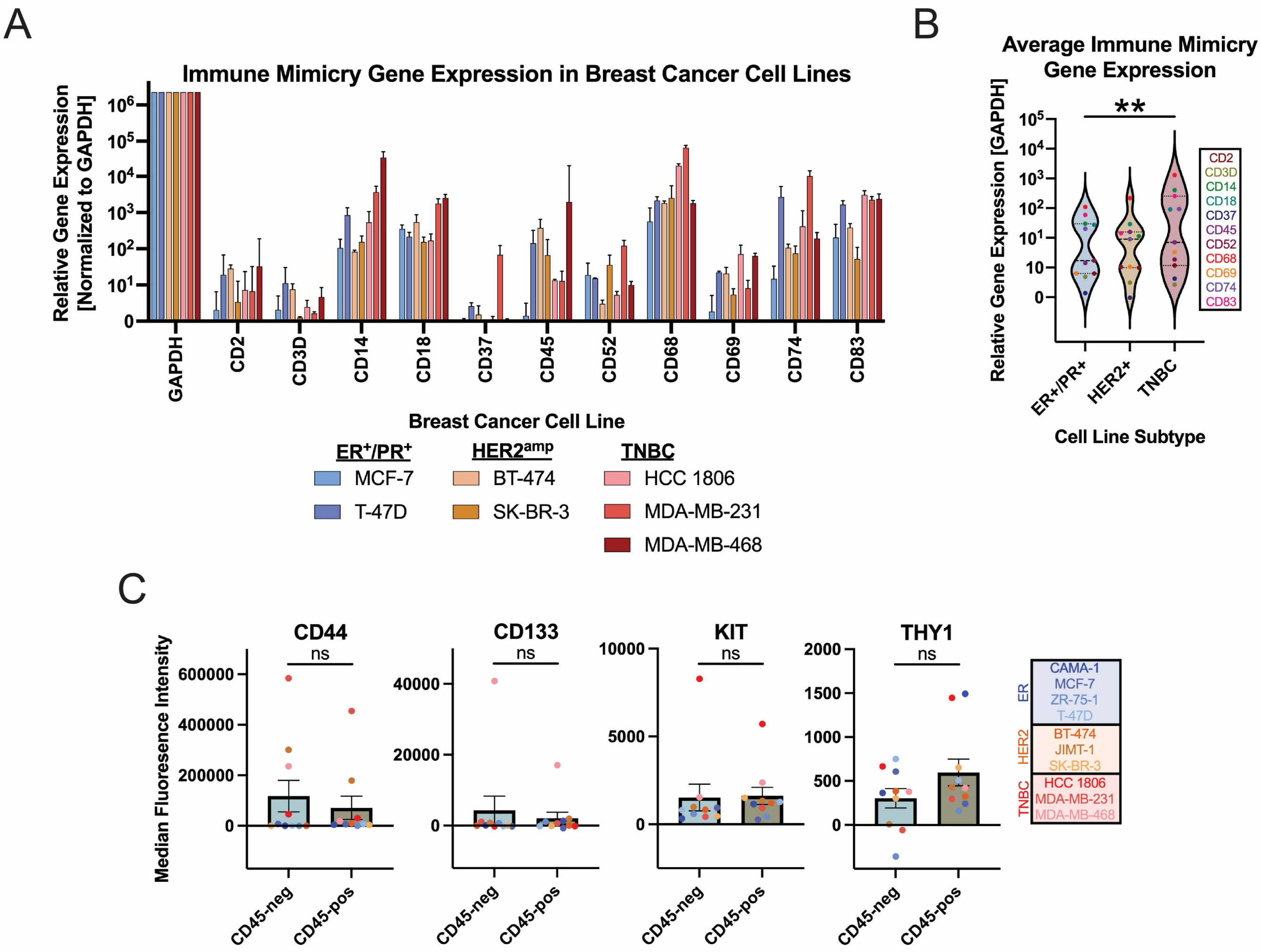


**Supplementary Figure 15**

**Supplementary Figure 15. Continued experimental validation of neoplastic immune mimicry in breast cancer cell lines.** (**A**) Evaluation of immune mimicry marker gene expression by qRT-PCR in a subset of breast cancer cell lines. (**B**) Triple-negative breast cancer cell lines express higher average levels of immune mimicry receptors at the RNA level when compared to hormone receptor-positive cell lines. (**C**) CD45-pos immune-mimicked cells do not significantly upregulate standard breast tumor-initiating markers like CD44, CD133, KIT, or THY1, supporting the notion that this is a novel cell state. ns = not significant, ** $p < 0.01$. One-way ANOVA with Dunnett correction (B) or Wilcoxon test (C). Data are means ± SEM.

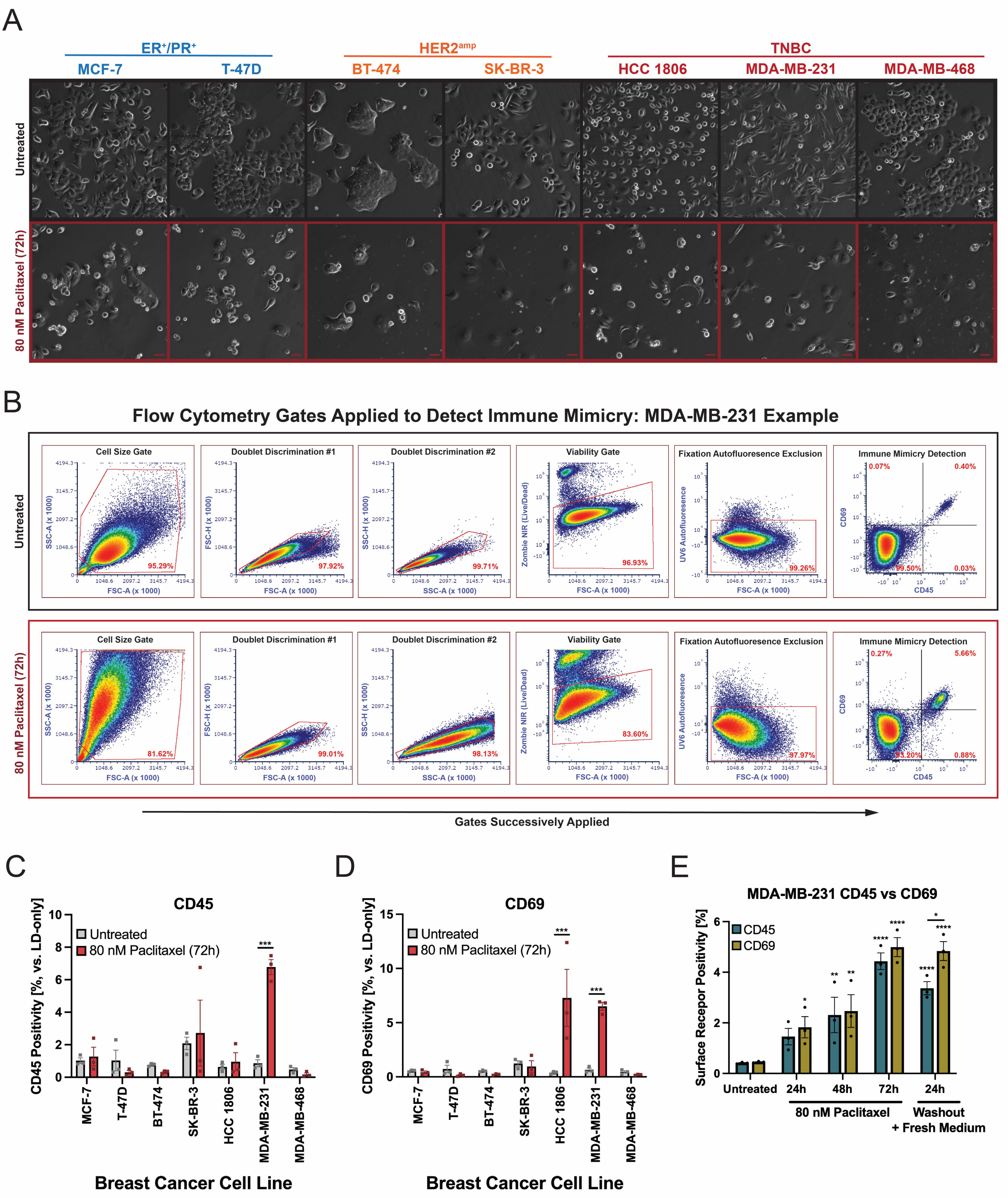


Supplementary Figure 16

**Supplementary Figure 16. CD45 and CD69 surface protein after paclitaxel treatment in a subset of breast cancer cell lines.** (**A**) Various breast cancer cell lines modeling different disease subtypes were subjected to 80 nM paclitaxel treatment for 72 hours. Surviving cells were subsequently collected for flow cytometric analysis. Scale bar, 25 μm. (**B**) Example of flow cytometry gates successively applied to detect neoplastic CD45 and CD69 on MDA-MB-231 cells in untreated vs paclitaxel treated culture conditions. (**C**) Significant induction of CD45 surface protein in response to paclitaxel was only observed for MDA-MB-231. Flow cytometric analysis represents all stains performed in biological triplicate. (**D**) Significant induction of CD69 surface protein after paclitaxel was observed for the triple-negative HCC 1806 and MDA-MB-231 cell lines. Flow cytometric analysis represents all stains performed in biological triplicate. (**E**) CD45 and CD69 surface staining in MDA-MB-231 cells along an 80 nM paclitaxel time-course reveals progressively increased CD45 and CD69 up to 72h, followed by CD45 and CD69 deviation after drug washout. Flow cytometric analysis represents all stains performed in biological triplicate. * $p < 0.05$, ** $p < 0.01$, *** $p < 0.001$, **** $p < 0.0001$. Two-way ANOVA with Šidák correction [(C), (D), and (E)]. Data are means ± SEM.

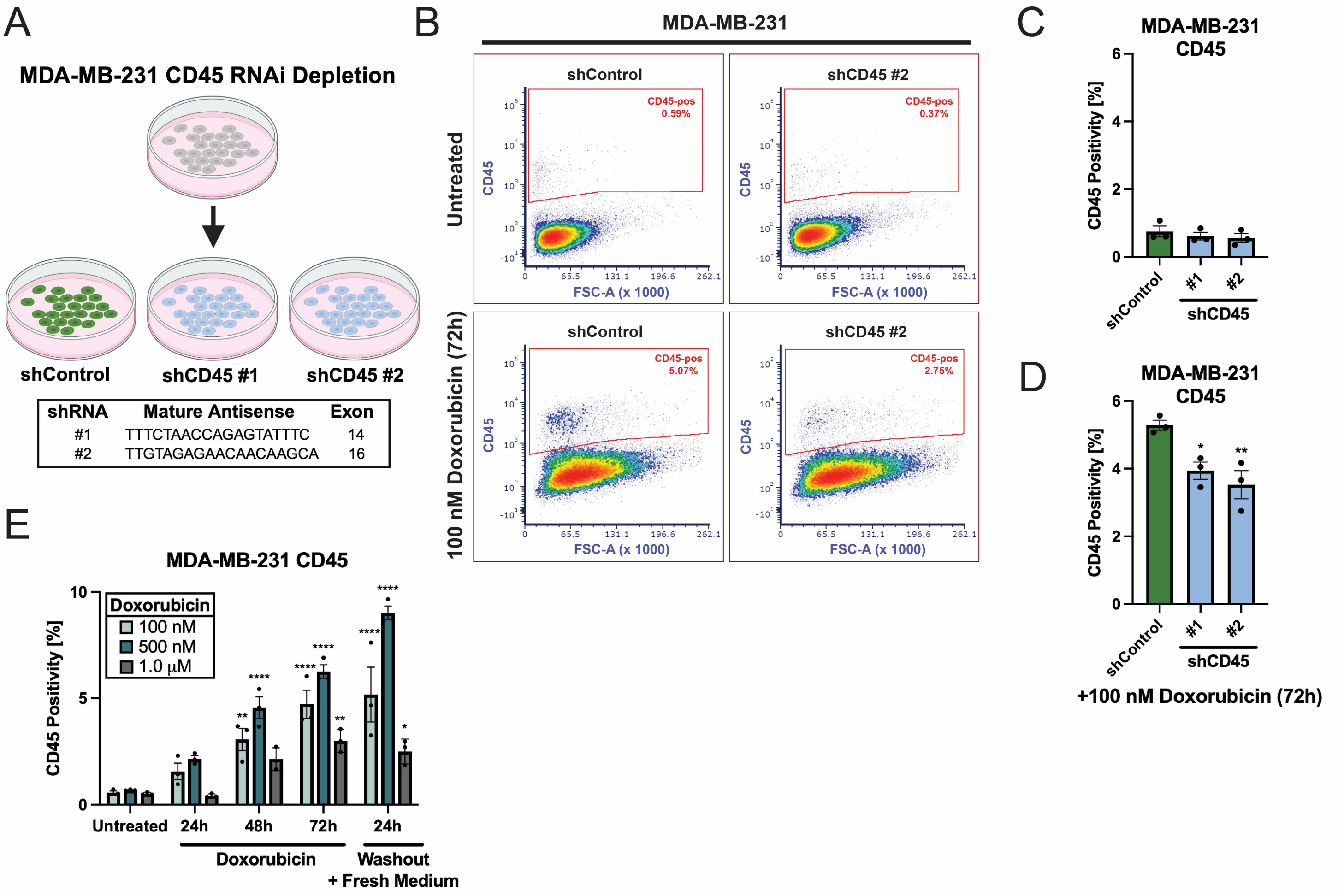


Supplementary Figure 17

**Supplemental Figure 17. CD45 depletion and expression time-course after doxorubicin in MDA-MB-231 cells.** (**A**) MDA-MB-231 cells were stably transduced with short hairpin RNA (shRNA) targeting CD45 (PTPRC). (**B-D**) Baseline CD45 expression in MDA-MB-231 cells is low, so doxorubicin treatment was used to elicit its expression, which revealed consistently lower levels in shCD45 lines compared to a control containing a non-silencing shRNA. (**E**) CD45 surface positivity measured along a doxorubicin time-course at different drug concentrations demonstrated reduced CD45 protein at a cytotoxic concentration of 1.0 μM. * $p < 0.05$; ** $p < 0.01$. One-way ANOVA with Dunnett post-test [(C) and (D)] or two-way ANOVA with Šidák correction [(E)]. Data are means ± SEM.

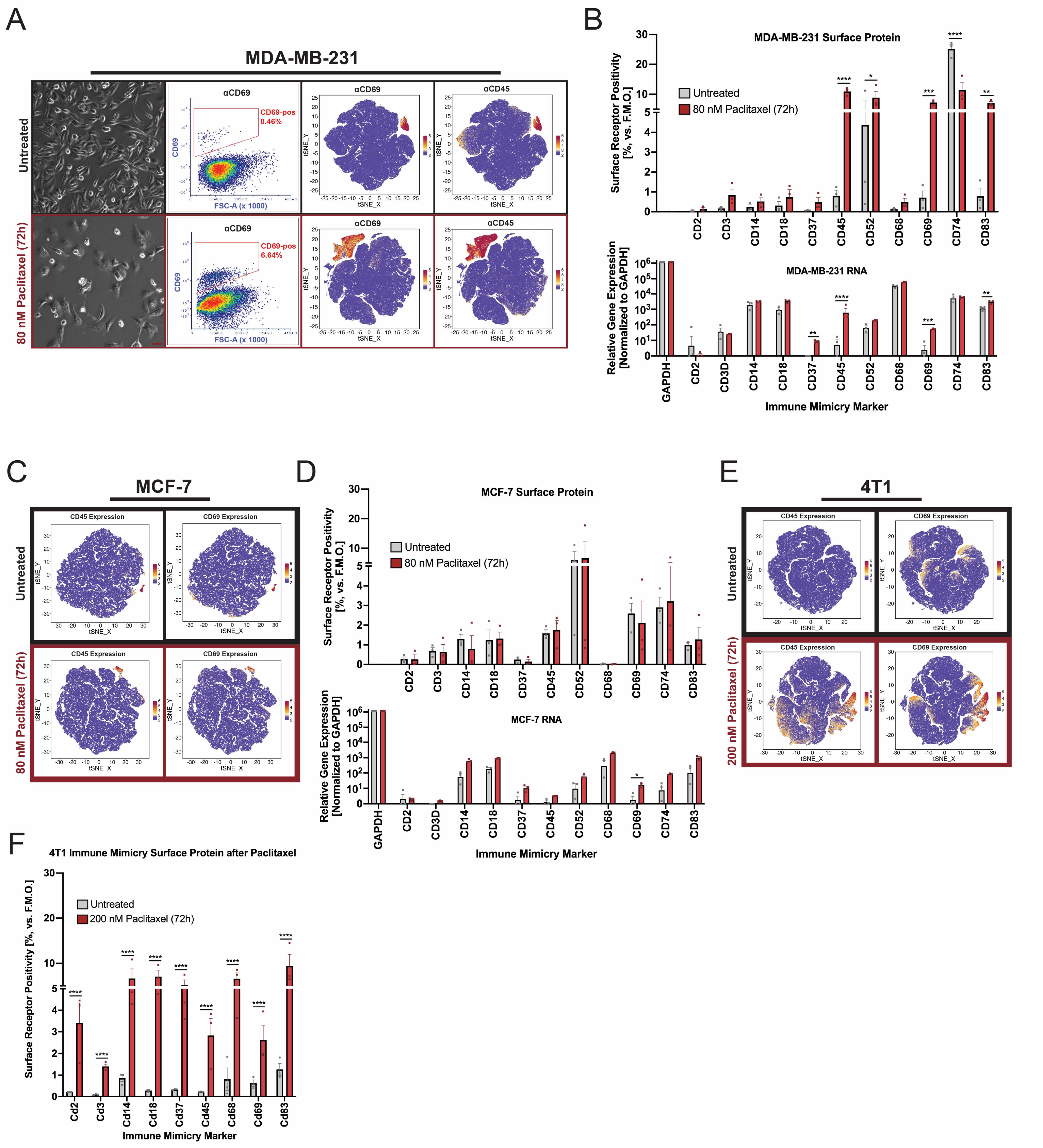


Supplementary Figure 18

**Supplementary Figure 18. Paclitaxel treatment elicits immune mimicry in aggressive cell lines.** (**A**) For aggressive and metastatic MDA-MB-231 cells, paclitaxel treatment leads to the expansion of an immune-like subpopulation expressing CD45 and CD69. Scale bar, 25 μm. (**B**) The upregulation of leukocyte surface receptors in MDA-MB-231 at the protein level is consistent with increased RNA expression. Flow cytometric analysis represents monostains; biological replicates were performed for both FACS and qRT-PCR. (**C**) For relatively non-metastatic MCF-7 cells, paclitaxel treatment does not expand the immune-like subpopulation expressing CD45 and CD69. (**D**) The lack of immune-mimicry induction in MCF-7 following paclitaxel is evidenced for surface protein as well as RNA. Flow cytometric analysis represents monostains; biological replicates were performed for both FACS and qRT-PCR. (**E-F**) For aggressive and metastatic murine mammary 4T1 cells, paclitaxel treatment leads to the widespread induction of several leukocyte surface receptors. Flow cytometric analysis represents monostains done in biological triplicate. * $p < 0.05$; ** $p < 0.01$; *** $p < 0.001$; **** $p < 0.0001$. Two-way ANOVA with Šidák correction [(B), (D) and (F)]. Data are means ± SEM.

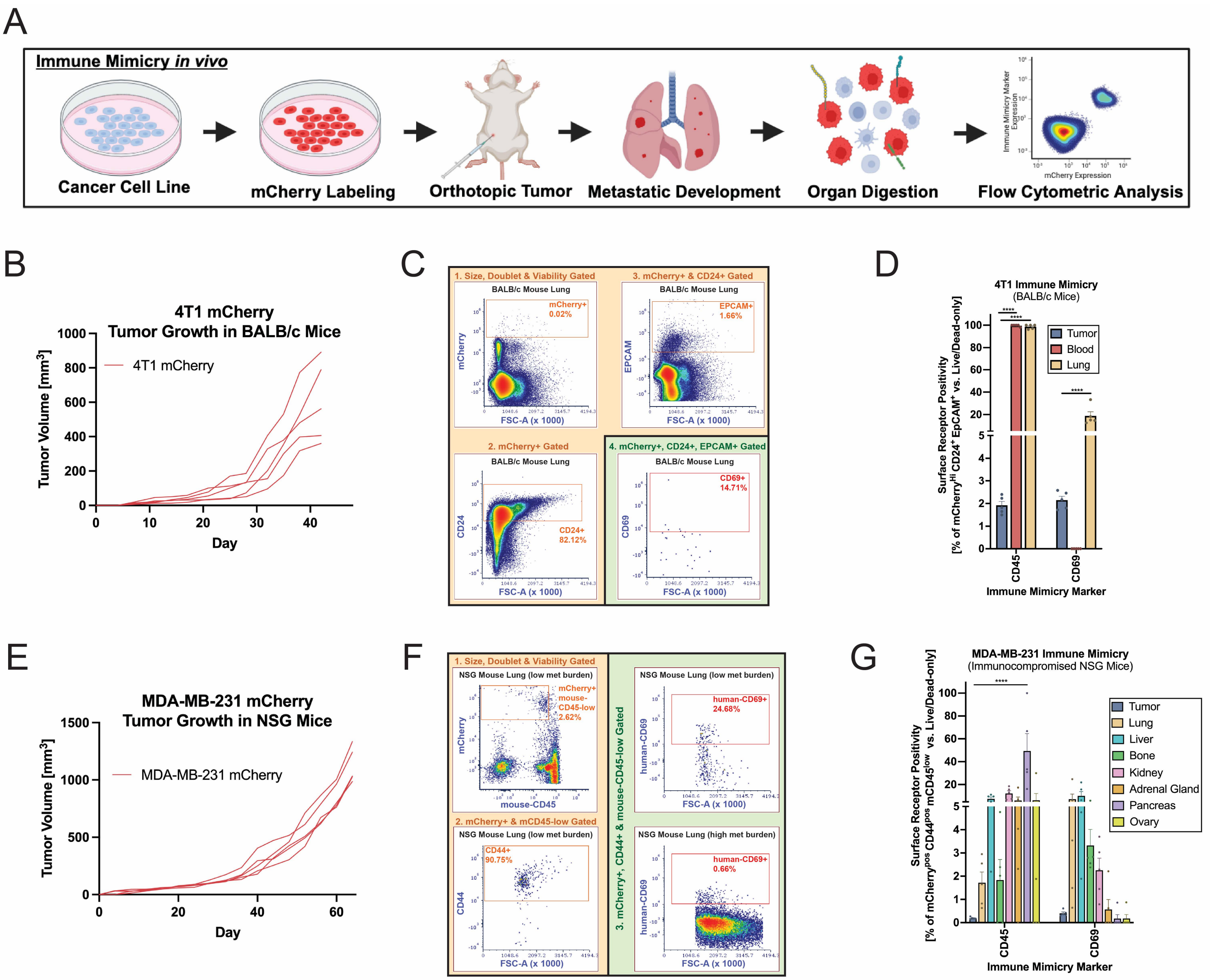


H

**MDA-MB-231 CD69 Positivity in NSG Lung Metastases**

| Mouse | Total MDA-MB-231 | CD69-pos MDA-MB-231 |
|---|---|---|
| Lung #1 | 0.08% | 24.68% |
| Lung #2 | 0.09% | 6.30% |
| Lung #3 | 0.23% | 3.54% |
| Lung #4 | 0.45% | 1.49% |
| Lung #5 | 2.45% | 0.66% |

Spearman Correlation: -1.00
$p = 0.0167$

**Supplementary Figure 19**

**Supplementary Figure 19. CD45 and CD69 are induced during 4T1 and MDA-MB-231 metastasis in mice.** (**A**) 4T1 and MDA-MB-231 cells were transduced with mCherry to facilitate sorting from metastatic tissues after orthotopic tumor growth in mice. (**B**) Tumor growth curves for 4T1 cells expressing mCherry after injection into BALB/c mice. (**C**) At endpoint, the tumor, blood, and lungs of mice were digested and the prospective flow cytometric analysis of 4T1 cells leveraged three redundant epithelial makers: mCherry, CD24, and EpCAM. (**D**) Metastatic 4T1 cells expressing mCherry, CD24, and EpCAM showed upregulation of CD45 in both the blood and lungs whereas CD69 was only upregulated in the lung. (**E**) Tumor growth curves for MDA-MB-231 cells expressing mCherry after injection into immunocompromised NSG mice. (**F**) Prospective identification of MDA-MB-231 cells from mouse tissue leveraged mCherry and CD44, which are two markers ubiquitously expressed in the cell line. These cells were also found to have lower levels of mouse-CD45 though amid suspicion that antibodies targeting mouse CD45 may have cross-reacted with the human epitope and vice versa. Examples of human CD69 staining in lungs with low versus high metastatic burden shown. (**G**) MDA-MB-231 cells ostensibly positive for CD45 and CD69 were retrieved from various metastatic tissues in immunocompromised NSG mice with (**H**) CD69 positivity being inversely correlated with lung metastatic burden. Antibodies used in this experiment targeted human CD45 and CD69 epitopes on MDA-MB-231 cells within dissociated mouse tissue. **** $p < 0.0001$. Two-way ANOVA with Šidák correction [(D) and (G)] or Spearman correlation (H). Data are means ± SEM.

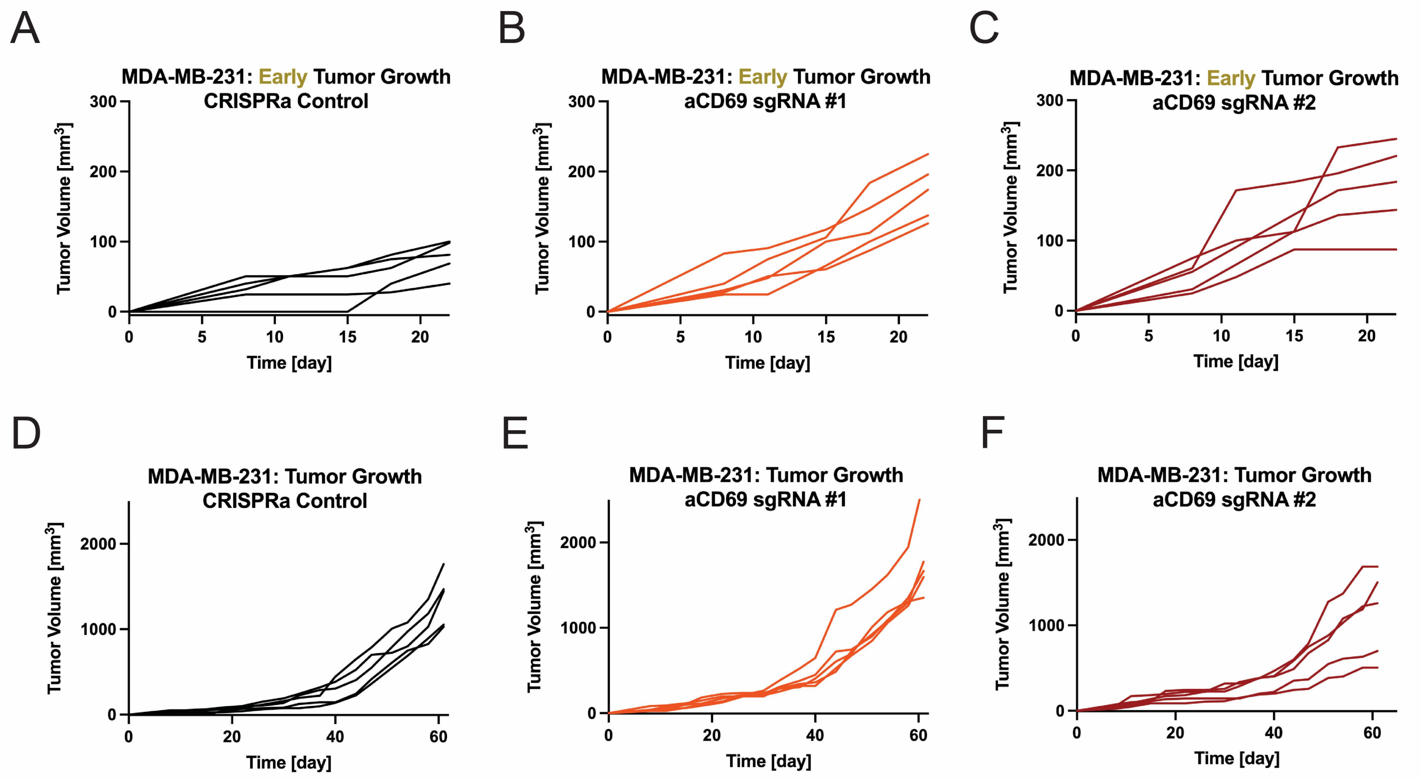


**Supplementary Figure 20**

**Supplementary Figure 20. Extended data for MDA-MB-231 CD69 CRISPR-activation experiments and the disruption of CD69 using CRISPR/Cas9.** (**A-F**) Spider plots showing the (**A-C**) early and (**D-F**) entire growth of individual tumors following the CRISPR-activation of CD69.

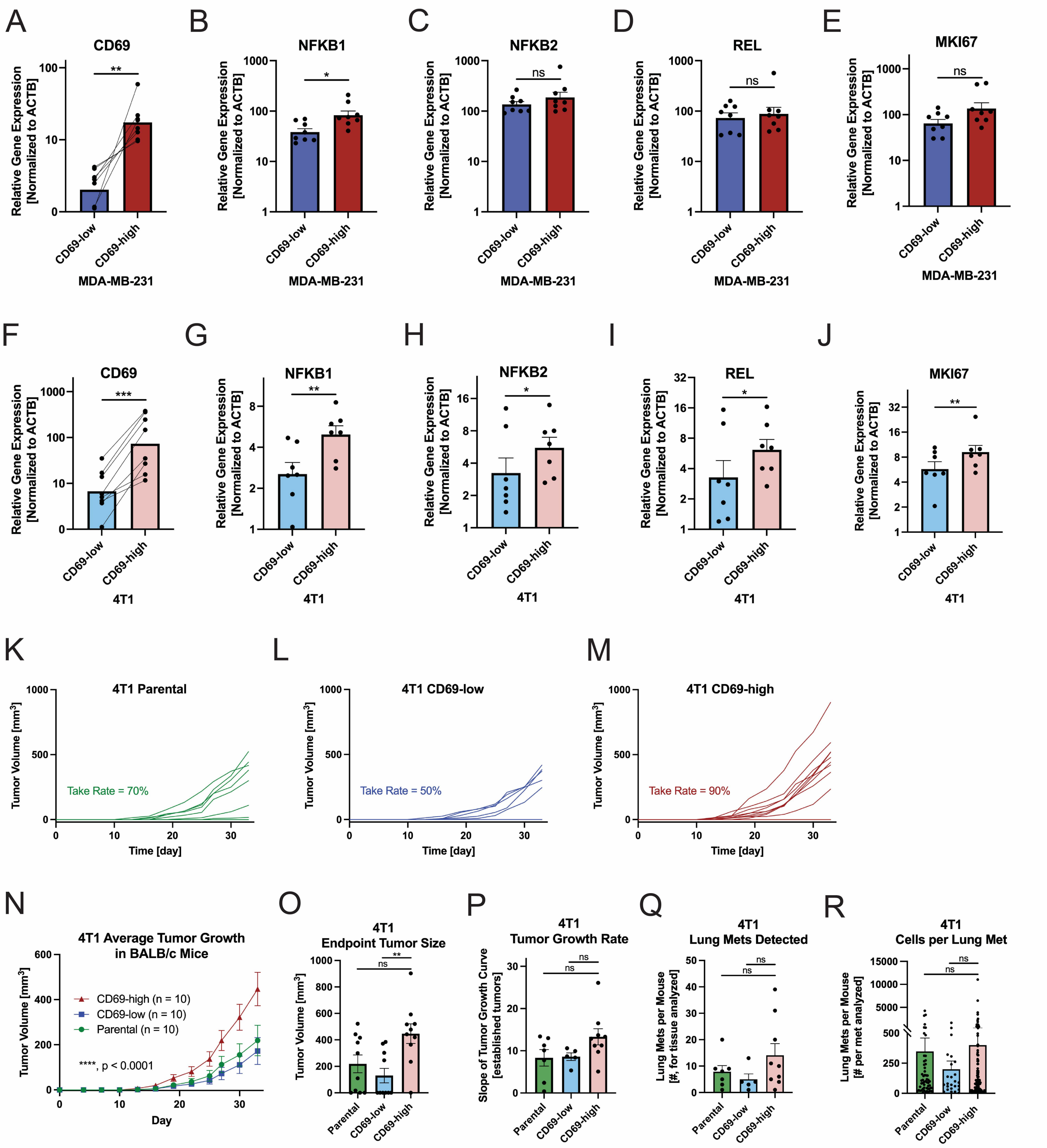


Supplementary Figure 21

**Supplementary Figure 21. Extended data for MDA-MB-231 and 4T1 CD69 magnetic sorting experiments.** (**A**-**E**) RNA expression of (**A**) CD69, (**B**-**D**) NFκB genes, and (**E**) MKI67 in sorted MDA-MB-231 cells. (**F**-**J**) RNA expression of (**F**) CD69, (**G**-**I**) NFκB genes, and (**J**) MKI67 in sorted 4T1 cells. (**K**-**M**) Spider plots exhibiting the growth of individual 4T1 tumors derived from the parental control versus CD69-low and CD69-high sorted subpopulations. (**N**) The outcome of this study summarized by (**N**) tumor growth, (**O**) endpoint tumor size, (**P**) the growth rate of established tumors, (**Q**) the number of lung metastases detected per mouse and (**R**) the cell size of individual lung metastases. ns = not significant, * $p < 0.05$; ** $p < 0.01$; *** $p <0.001$; **** $p < 0.0001$. Student's t-test [(A-J)], two-way ANOVA, or one-way ANOVA with Tukey correction [(O-R)]. Data are means ± SEM.

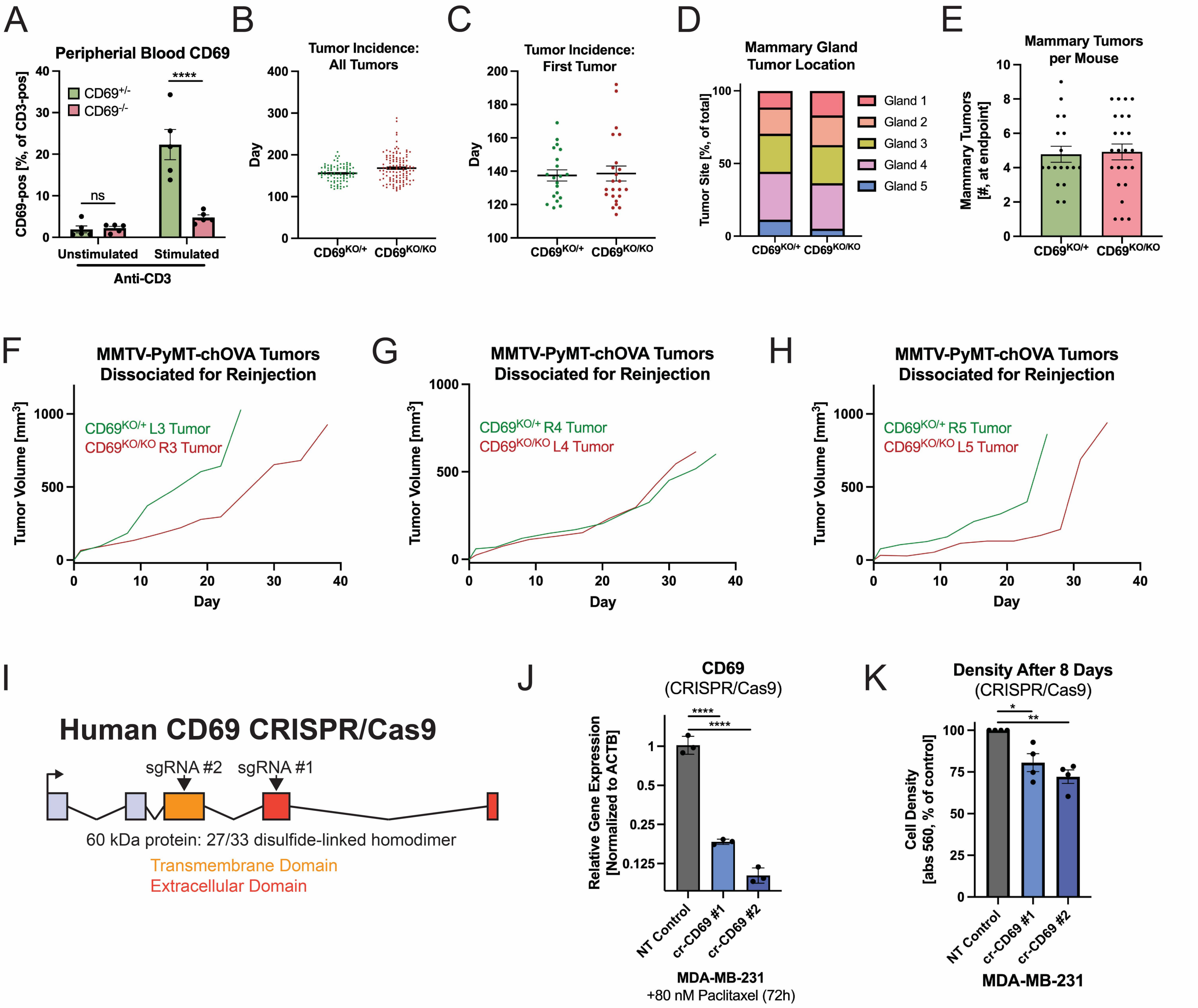


**Supplementary Figure 22**

**Supplementary Figure 22. Extended data for MMTV-PyMT-CD69$^{KO/KO}$ experiments.** (**A**) The CD69 protein is reduced in CD69$^{KO/KO}$ mice demonstrated here on CD3-pos cells by flow cytometry following anti-CD3 stimulation of peripheral blood. (**B**-**C**) There is no statistical difference in the detection time of spontaneous tumors for the (**B**) first tumor or (**C**) all tumors in MMTV-PyMT-CD69$^{KO/+}$ and MMTV-PyMT-CD69$^{KO/KO}$ mice. (**D**) The distribution of mammary tumors and (**E**) the number developed in MMTV-PyMT-CD69$^{KO/+}$ and MMTV-PyMT-CD69$^{KO/KO}$ mice is similar. (**F**-**H**) Tumors attaining similar endpoint sizes were dissociated for reinjection back into CD69$^{KO/+}$ mice. (**I**) CD69 CRISPR/Cas9 sgRNAs targeting the protein's extracellular or transmembrane domain were transduced into MDA-MB-231 along with constitutively active Cas9. (**J**) MDA-MB-231 cells with disrupted CD69 have reduced RNA expression following paclitaxel treatment, which was done to compare CD69 induction capacity. (**K**) MDA-MB-231 cells harboring disrupted CD69 achieve lower density following growth *in vitro*. * $p < 0.05$; ** $p < 0.01$; **** $p < 0.0001$. Two-way ANOVA with Šidák correction [(A)] or one-way ANOVA with Tukey correction [(J) and (K)]. Data are means ± SEM.

**Table S1. Clinical information for patients in the PANNTHR Trial at the Oregon Health & Science University.**

| Patient | Age at Enrollment | Histology | Histology (ICD-O-3) | Primary Site | Primary Site (ICD-O-3) |
|---|---|---|---|---|---|
| Patient 1 | 28 | Infiltrating duct carcinoma, NOS | 8500 | Lower-outer quadrant of breast | C50.5 |
| Patient 2 | 52 | Infiltrating duct carcinoma, NOS | 8500 | Overlapping lesion of breast | C50.8 |
| Patient 3 | 53 | Infiltrating duct carcinoma, NOS | 8500 | Upper breast | C50.8 |
| Patient 4 | 60 | Infiltrating duct carcinoma, NOS | 8500 | Upper-outer quadrant of breast | C50.4 |
| Patient 5 | 62 | Infiltrating duct carcinoma, NOS | 8500 | Upper-outer quadrant of breast | C50.4 |
| Patient 6 | 39 | Infiltrating duct carcinoma, NOS | 8500 | Overlapping lesion of breast | C50.8 |
| Patient 7 | 59 | Infiltrating duct carcinoma, NOS | 8500 | Lower-outer quadrant of breast | C50.5 |

**Table S1 (cont.). Clinical information for patients in the PANNTHR Trial at the Oregon Health & Science University.**

| Patient | Laterality | PANNTHR Study Treatment | Other Neoadjuvant Treatments |
|---|---|---|---|
| Patient 1 | Left | Niraparib, Abemaciclib | Goserelin, Anastrozole, Leuprolide, Cyclophosphamide, Docetaxel |
| Patient 2 | Right | Niraparib, Abemaciclib | Tamoxifen, Goserelin, Letrozole |
| Patient 3 | Right | Niraparib, Abemaciclib | Letrozole |
| Patient 4 | Left | Niraparib, Abemaciclib | Letrozole |
| Patient 5 | Left | Niraparib, Abemaciclib | Letrozole, Carboplatin, Docetaxel, Pertuzumab, Trastuzumab |
| Patient 6 | Right | Niraparib, Abemaciclib | Tamoxifen, Cyclophosphamide, Docetaxel |
| Patient 7 | Right | Niraparib, Abemaciclib | Letrozole, Trastuzumab, Pertuzumab |

**Table S1 (cont.). Clinical information for patients in the PANNTHR Trial at the Oregon Health & Science University.**

| Patient | Follow-up Days after Surgery | Recurrence at Last Follow Up? |
|---|---|---|
| Patient 1 | 544 | No |
| Patient 2 | 652 | No |
| Patient 3 | 755 | No |
| Patient 4 | 791 | No |
| Patient 5 | 336 | No |
| Patient 6 | 783 | No |
| Patient 7 | 518 | No |

**Table S2. Reagents used for multiplexed immunohistochemistry staining.**

| Staining Order | Marker | Antibody Clone | Vendor |
|---|---|---|---|
| 1 | Hematoxylin & Eosin | | Hematoxylin: Vector, Eosin: Fisher |
| 2 | CD69 | Polyclonal | Sigma |
| 3 | CD45 | H130 | Fisher |
| 4 | CD3 | SP7 | Fisher |
| 5 | CD20 | L26 | Abcam |
| 6 | CD68 | PG-M1 | Abcam |
| 7 | Vimentin | 280618 | R&D Systems |
| 8 | CD11b | EPR1344 | Abcam |
| 9 | CD163 | 10D6 | ThermoFisher Scientific |
| 10 | CD31 | Polyclonal | Abcam |
| 11 | CD8 | C8/144B | Fisher |
| 12 | KRT19 | Polyclonal | Sigma |
| 13 | PanCK | AE1/AE3//5D3 | Abcam |
| 14 | KI67 | SP6 | Sigma |
| 15 | H3/NUCA | H3: D1H2, NucA: NM106 | H3: Cell Signaling Technologies, NucA: Abcam |
| 16 | Hematoxylin | | Vector |

**Table S2 (cont.). Reagents used for multiplexed immunohistochemistry staining.**

| Marker | Catalog # | RRID |
|---|---|---|
| Hematoxylin & Eosin | Hematoxylin: H-3401-500, Eosin: 22-110-637 | |
| CD69 | HPA050525 | RRID:AB_2681157 |
| CD45 | 50-126-09 | RRID:AB_467274 |
| CD3 | RM9107S | RRID:AB_149924 |
| CD20 | ab9475 | RRID:AB_307267 |
| CD68 | ab783 | RRID:AB_306119 |
| Vimentin | MAB2105 | RRID:AB_2241653 |
| CD11b | ab133357 | RRID:AB_2650514 |
| CD163 | MA5-11458 | RRID:AB_10982556 |
| CD31 | Ab28364 | RRID:AB_726362 |
| CD8 | PIMA513473 | RRID:AB_11000353 |
| KRT19 | HPA002465 | RRID:AB_1079179 |
| PanCK | ab86734 | RRID:AB_10674321 |
| KI67 | 275R-17 | RRID:AB_1158035 |
| H3/NUCA | H3: 4499L NucA: ab215396 | H3: RRID:AB_10544537, NucA: RRID: AB_3677433 |
| Hematoxylin | H-3401-500 | |

**Table S3. Cell lines used in this study.**

| Cell Line | Supplier | Catalog | Passage |
|---|---|---|---|
| CAMA-1 | ATCC | HTB-21 | Passage 52 |
| MCF-7 | ATCC | HTB-22 | Passage 22 |
| T-47D | ATCC | HTB-133 | Passage 4 |
| ZR-75-1 | ATCC | CRL-1500 | Passage 12 |
| BT-474 | ATCC | HTB-20 | Passage 9 |
| JIMT-1 | DSMZ | ACC 589 | Passage 25 |
| SK-BR-3 | ATCC | HTB-30 | Passage 11 |
| HCC 1806 | ATCC | CRL-2335 | Passage 6 |
| MDA-MB-231 | ATCC | HTB-26 | Passage 8 |
| MDA-MB-468 | ATCC | HTB-132 | Passage 15 |
| 4T1 | ATCC | CRL-2539 | Passage 10 |
| PyMT chOVA | Zena Werb Lab | | Passage 14 |

**Table S3 (cont.). Cell lines used in this study.**

| Cell Line | Mycoplasma Free? | Mycoplasma Testing | RRID |
|---|---|---|---|
| CAMA-1 | Yes | In-house PCR | RRID:CVCL_1115 |
| MCF-7 | Yes | Charles River | RRID:CVCL_0031 |
| T-47D | Yes | In-house PCR | RRID:CVCL_0553 |
| ZR-75-1 | Yes | In-house PCR | RRID:CVCL_0588 |
| BT-474 | Yes | In-house PCR | RRID:CVCL_0179 |
| JIMT-1 | Yes | In-house PCR | RRID:CVCL_2077 |
| SK-BR-3 | Yes | In-house PCR | RRID:CVCL_0033 |
| HCC 1806 | Yes | Charles River | RRID:CVCL_1258 |
| MDA-MB-231 | Yes | Charles River | RRID:CVCL_0062 |
| MDA-MB-468 | Yes | In-house PCR | RRID:CVCL_0419 |
| 4T1 | Yes | Charles River | RRID:CVCL_0125 |
| PyMT chOVA | Yes | Charles River | |

**Table S4. Anti-human antibodies used for flow cytometric analyses.**

| Marker | Antibody Clone | Fluorophore | Company | Catalog | RRID |
|---|---|---|---|---|---|
| CD2 | TS1/8 | PE/Cy5 | Biolegend | 300209 | RRID:AB_314033 |
| CD3 | SK7 | BV 750 | Biolegend | 344845 | RRID:AB_2734352 |
| CD14 | M5E2 | BV 711 | Biolegend | 301837 | RRID:AB_11218986 |
| CD18 | TS1/18 | PE/Dazzle 594 | Biolegend | 302127 | RRID:AB_2750396 |
| CD24 | ML5 | PerCP | Biolegend | 311113 | RRID:AB_2561283 |
| CD37 | M-B371 | FITC | Biolegend | 356303 | RRID:AB_2561836 |
| CD44 | IM7 | Pac Blue | Biolegend | 103020 | RRID:AB_493683 |
| CD45 | HI30 | AF 700 | Biolegend | 304023 | RRID:AB_493760 |
| CD52 | HL186 | PerCP/Cy5.5 | Biolegend | 316009 | RRID:AB_2650806 |
| CD68 | Y1/82A | BV 785 | Biolegend | 333825 | RRID:AB_2800879 |
| CD69 | FN50 | PE/Cy7 | Biolegend | 310912 | RRID:AB_314847 |
| CD74 | LN2 | PE | Biolegend | 326807 | RRID:AB_2229059 |
| CD83 | HB15e | APC/Fire 750 | Biolegend | 305331 | RRID:AB_2650746 |
| CD133 | S16015F | BV 421 | Biolegend | 393907 | RRID:AB_2832742 |
| KIT (CD117) | 104D2 | BV 605 | Biolegend | 313217 | RRID:AB_2562024 |
| THY1 (CD90) | 5E10 | BV 650 | Biolegend | 328143 | RRID:AB_2734319 |
| Ki67 | Ki-67 | Pac Blue | Biolegend | 350511 | RRID:AB_10895904 |
| Live/Dead | | Zombie NIR | Biolegend | 423106 | |
| CD69 | FN50 | APC | Biolegend | 985206 | RRID:AB_2922660 |
| Live/Dead | | Zombie Violet | Biolegend | 423114 | |

**Table S5. Anti-murine antibodies used for flow cytometric analyses.**

| Marker | Antibody Clone | Fluorophore | Company | Catalog | RRID |
|---|---|---|---|---|---|
| CD2 | RM2-5 | PerCP/Cy5.5 | Biolegend | 100115 | RRID:AB_2563501 |
| CD3 | 17A2 | BV 750 | Biolegend | 100249 | RRID:AB_2734148 |
| CD14 | Sa14-2 | APC/Fire 750 | Biolegend | 123331 | RRID:AB_2734179 |
| CD18 | M18/2 | FITC | Biolegend | 101405 | RRID:AB_312814 |
| CD24 | M1/69 | AF 700 | Biolegend | 101835 | RRID:AB_2566729 |
| CD37 | Duno85 | PE | Biolegend | 146203 | RRID:AB_2562285 |
| CD44 | IM7 | Pac Blue | Biolegend | 103020 | RRID:AB_493683 |
| CD45 | 30-F11 | BV 650 | Biolegend | 103151 | RRID:AB_2565884 |
| CD68 | FA-11 | BV 711 | Biolegend | 137029 | RRID:AB_2783098 |
| CD69 | H1.2F3 | APC | Biolegend | 104514 | RRID:AB_492843 |
| CD83 | Michel-19 | PE/Cy7 | Biolegend | 121517 | RRID:AB_2566123 |
| CD133 | 315-2C11 | BV 421 | Biolegend | 141213 | RRID:AB_2566011 |
| EpCAM (CD326) | G8.8 | BV 510 | Biolegend | 118231 | RRID:AB_2632774 |
| KIT (CD117) | AKC2 | BV 785 | Biolegend | 135138 | RRID:AB_2734197 |
| THY1 (CD90) | 30-H12 | PE/Cy5 | Biolegend | 105314 | RRID:AB_313185 |
| Ki67 | B56 | BV 480 | BD Biosciences | 566109 | RRID:AB_2739511 |
| CD45 | 30-F11 | PerCP | Biolegend | 103130 | RRID:AB_893339 |
| Live/dead | | Zombie NIR | Biolegend | 423106 | |

**Table S6. qRT-PCR primers used in this study.**

| Primer | Species | Sequence |
|---|---|---|
| hACTB_FWD | Human | CATGTACGTTGCTATCCAGGC |
| hACTB_REV | Human | CTCCTTAATGTCACGCACGAT |
| hGAPDH_met_FWD | Human | TCAAGGCTGAGAACGGGAAG |
| hGAPDH_met_REV | Human | CGCCCCACTTGATTTTGGAG |
| hCD2_FWD | Human | TCAAGAGAGGGTCTCAAAACCA |
| hCD2_REV | Human | CCATTCATTACCTCACAGGTCAG |
| hCD3D_FWD | Human | ACTGGCTACCCTTCTCTCG |
| hCD3D_REV | Human | CCGTTCCCTCTACCCATGTGA |
| hCD14_FWD | Human | ACGCCAGAACCTTGTGAGC |
| hCD14_REV | Human | GCATGGATCTCCACCTCTACTG |
| hCD18_FWD | Human | TGCGTCCTCTCTCAGGAGTG |
| hCD18_REV | Human | GGTCCATGATGTCGTCAGCC |
| hCD37_FWD | Human | GCTGGGACTATGTGCAGTTCC |
| hCD37_REV | Human | ACCCGTTACCTCTCAGGATGA |
| hCD45_FWD | Human | ACCACAAGTTTACTAACGCAAGT |
| hCD45_REV | Human | TTTGAGGGGGATTCCAGGTAAT |
| hCD52_FWD | Human | TCTTCCTCCTACTCACCATCAG |
| hCD52_REV | Human | CCTCCGCTTATGTTGCTGGA |
| hCD68_FWD | Human | TGGGGCAGAGCTTCAGTTG |
| hCD68_REV | Human | TGGGGCAGGAGAAACTTTGC |
| hCD69_FWD | Human | ATTGTCCAGGCCAATACACATT |
| hCD69_REV | Human | CCTCTCTACCTGCGTATCGTTTT |
| hCD74_FWD | Human | GACGAGAACGGCAACTATCTG |
| hCD74_REV | Human | GTTGGGGAAGACACACCAGC |
| hCD83_FWD | Human | AAGGGGCAAAATGGTTCTTTCG |
| hCD83_REV | Human | GCACCTGTATGTCCCCGAG |
| hCD69_upstream_FWD | Human | GTAGCTTGACTTGACCTGAGATT |
| hCD69_upstream_REV | Human | ATGGTGATGAAGACCACATTCA |
| hNFKB1_FWD | Human | AACAGAGAGGATTTCGTTTCCG |
| hNFKB1_REV | Human | TTTGACCTGAGGGTAAGACTTCT |

| | | |
|---|---|---|
| hNFKB2_FWD | Human | ATGGAGAGTTGCTACAACCCA |
| hNFKB2_REV | Human | CTGTTCCACGATCACCAGGTA |
| hREL_FWD | Human | GCAGAGGGGAATGCGTTTTAG |
| hREL_REV | Human | AGAAGGGTATGTTCGGTTGTTG |
| hMKI67_FWD | Human | GCCTGCTCGACCCTACAGA |
| hMKI67_REV | Human | GCTTGTCAACTGCGGTTGC |
| mActb_FWD | Mouse | GGCTGTATTCCCCTCCATCG |
| mActb_REV | Mouse | CCAGTTGGTAACAATGCCATGT |
| mGapdh_FWD | Mouse | AGGTCGGTGTGAACGGATTTG |
| mGapdh_REV | Mouse | TGTAGACCATGTAGTTGAGGTCA |
| mCd69_FWD | Mouse | CCCTTGGGCTGTGTTAATAGTG |
| mCd69_REV | Mouse | AACTTCTCGTACAAGCCTGGG |
| mCd45_FWD | Mouse | GTTTTCGCTACATGACTGCACA |
| mCd45_REV | Mouse | AGGTTGTCCAACTGACATCTTTC |
| mNfkb1_Fwd | Mouse | ATGGCAGACGATGATCCCTAC |
| mNfkb1_Rev | Mouse | TGTTGACAGTGGTATTTCTGGTG |
| mNfkb2_Fwd | Mouse | GGCCGGAAGACCTATCCTACT |
| mNfkb2_Rev | Mouse | CTACAGACACAGCGCACACT |
| mRel_Fwd | Mouse | AGAGGGGAATGCGGTTTAGAT |
| mRel_Rev | Mouse | TTCTGGTCCAAATTCTGCTTCAT |
| mMki67_Fwd | Mouse | ATCATTGACCGCTCCTTTAGGT |
| mMki67_Rev | Mouse | GCTCGCCTTGATGGTTCCT |

**Table S7. Oligonucleotides used for cloning CRISPR-a and CRISPR/Cas9 plasmids.**

| sgRNA Oligo | Sequence |
|---|---|
| hCD69_CRISPRa_sg1_top | CACCGTGTGTTGTTGTGGTGAAGTA |
| hCD69_CRISPRa_sg1_bottom | AAACTACTTCACCACAACAACACAC |
| hCD69_CRISPRa_sg2_top | CACCGCTATACATTGTCTGGTCCAC |
| hCD69_CRISPRa_sg2_bottom | AAACGTGGACCAGACAATGTATAGC |
| hCD69_CRISPRcas9_sg1_top | CACCGAACTTTCTAAAACGATACGC |
| hCD69_CRISPRcas9_sg1_bottom | AAACGCGTATCGTTTTAGAAAGTTC |
| hCD69_CRISPRcas9_sg2_top | CACCGGTGGGCCAATACAATTGTCC |
| hCD69_CRISPRcas9_sg2_bottom | AAACGGACAATTGTATTGGCCCACC |

**Table S8. Plasmids used in this study.**

| Plasmid | Vendor | Catalog | RRID |
|---|---|---|---|
| pHAGE TRE dCas9-VP64 | Addgene | 50916 | RRID:Addgene_50916 |
| lenti sgRNA(MS2)_Puro backbone | Addgene | 73795 | RRID:Addgene_73795 |
| LentiMPH v2 | Addgene | 89308 | RRID:Addgene_89308 |
| pLV-eGFP | Addgene | 36083 | RRID:Addgene_36083 |
| lentiCRISPR v2 | Addgene | 52961 | RRID:Addgene_52961 |
| pLV-mCherry | Addgene | 36084 | RRID:Addgene_36084 |
